# Dissecting functional dynamics of fecal-derived *in vitro* stable microbial communities with metaproteomics

**DOI:** 10.64898/2026.10.05.754657

**Authors:** Angela Wang, Emily EF Fekete, Marybeth Creskey, Zhibin Ning, Janice Mayne, Kai Cheng, Qing Wu, Vanessa M. D’Costa, Xuguang Li, Daniel Figeys, Xu Zhang

**Author notes:** Correspondence to (XZ), (DF), and (SL).

## Abstract

Fecal-derived *in vitro* microbial communities represent promising models for investigating gut microbiome function and facilitating translational applications. However, the factors governing community establishment and the underlying mechanisms remain incompletely understood. We employed large-scale quantitative metaproteomics to investigate the functional and community dynamics of human fecal microbiomes cultured in three media over 25 serial passages. Our results demonstrated that both culture medium and donor-specific microbiome characteristics significantly influence community establishment. Taxon-specific functional analyses revealed nutrient availability in the medium and the microbial stress responses as key determinants of *in vitro* community establishment. Notably, the metabolism of mucin glycans in RapidAIM medium contributed to the maintenance of *Bacteroidaceae* species that were poorly supported in other media, resulting in an *in vitro* community that most closely resembled the uncultured fecal inoculum. Overall, these findings provide mechanistic insights into *in vitro* gut microbial community establishment, supporting further optimization and application of fecal-derived *in vitro* microbiome models.

## Introduction

Often referred to as a “forgotten organ”, the human gut microbiome has profound influences on host physiology through synergistic interactions with the host to regulate digestion, nutrient absorption, metabolic function, and immune responses [1, 2]. The disruption of gut microbial homeostasis, or dysbiosis, leads to aberrant host-microbiome interactions and the onset of various diseases, including the gastrointestinal, cardiovascular diseases, and disorders in immune and nervous systems [3, 4]. Despite that mechanisms underlying the roles of microbiome in many diseases remain to be fully elucidated, targeting the gut microbiome for disease treatment has emerged as an important therapeutic strategy. Major milestones were achieved recently with the authorization of the first fecal microbiota transplant (FMT) product *Rebyota* in 2022 and the first oral capsule FMT therapeutics *Vowst* in 2023 in the United States for the prevention of recurrent *Clostridioides difficile* infection (rCDI) due to their clinical benefits [5–8]. However, the wide application of FMT products remains challenging due to the high complexity and inter-individual differences of donor fecal materials. Additionally, there is a growing need for suitable preclinical microbiome models that are stable, scalable, and reproducible. Such models are essential for applications, such as high-throughput drug screening and functional assessment of microbiome products [9].

In recent years, several *in vitro* models have been developed to enable functional microbiome studies and application. RapidAIM (rapid assay of an individual’s microbiome) is a high-throughput batch-culturing based platform [10, 11], which has been widely used for assessing microbiome interactions with xenobiotics, such as drugs [11, 12], resistant starches [13, 14], and natural compounds [15]. Continuous culturing platforms, such as SHIME (simulator of human intestinal microbial ecosystem), a multicompartment dynamic simulator for the entire gastrointestinal tract [16], have been used in various research fields, including food, nutrition and drug development [17]. There are also several synthetic microbial communities, such as hComs [18], providing compositionally defined and scalable systems for more controlled mechanistic investigations. Another scalable microbiome system that captures a greater degree of complexity of the gut microbiome is the fecal-derived stable *in vitro* microbial community (SIC). It has been reported that fecal-derived SICs can be achieved through serial passaging of stool microbiomes in suitable culture media, and when colonized in germ-free mice, the SICs reassembled natural microbiome composition and host proteomic responses [19]. The *in vitro* co-culture assay with SIC and pathogens successfully predicted the pathogen-colonization resistance of its precedent natural microbiome *in vivo* [19], highlighting its high functional and translational potential. However, to fully harness the translational potential of fecal-derived SICs, comprehensive characterization of the community and functional dynamics as well as the factors influencing their establishment is still needed.

Previous work with SICs generated with Brain-Heart Infusion (BHI) medium supplemented with different carbon sources suggested that nutrient availability is a key determinant of SIC assembly, driven by an early-resource utilization dynamic in which fast-growing taxa rapidly consume available substrates and subsequently dominate community structure [20]. However, the molecular mechanisms and functional responses underlying these compositional changes remain poorly understood. In particular, how microbial pathways are altered in different nutritional conditions has yet to be elucidated, which would benefit from multi-omics approaches. Metaproteomics, which directly measures the expressed proteins within the microbial community, is very well suited for functional profiling and mechanistic investigations [21, 22]. In particular, recent advances and application of state-of-the-art mass spectrometers (MS), sophisticated bioinformatic tools, and efficient data-independent acquisition (DIA) MS workflows in metaproteomics [23–25], has made it a powerful tool for microbiome investigations. In this study, we performed a quantitative DIA-MS metaproteomics to investigate the SIC establishments in multiple culture media with distinct nutrient compositions, including those that were designed to better maintain gut microbiome functionalities. Using six donor-derived fecal microbiomes cultured over 25 serial passages and sampled at representative time points for metaproteomic analysis, this study profiled 1008 metaproteomic samples, together with corresponding microbial growth and supernatant pH measurements (Figure 1A). The resulting large-scale metaproteomic data enables a systematic investigation of how nutrient availability and donor-specific microbial composition influence the establishment, functional adaptation, and stability of gut microbial communities *in vitro*, providing mechanistic insights that can advance their further development and broader applications.

**Figure 1.**
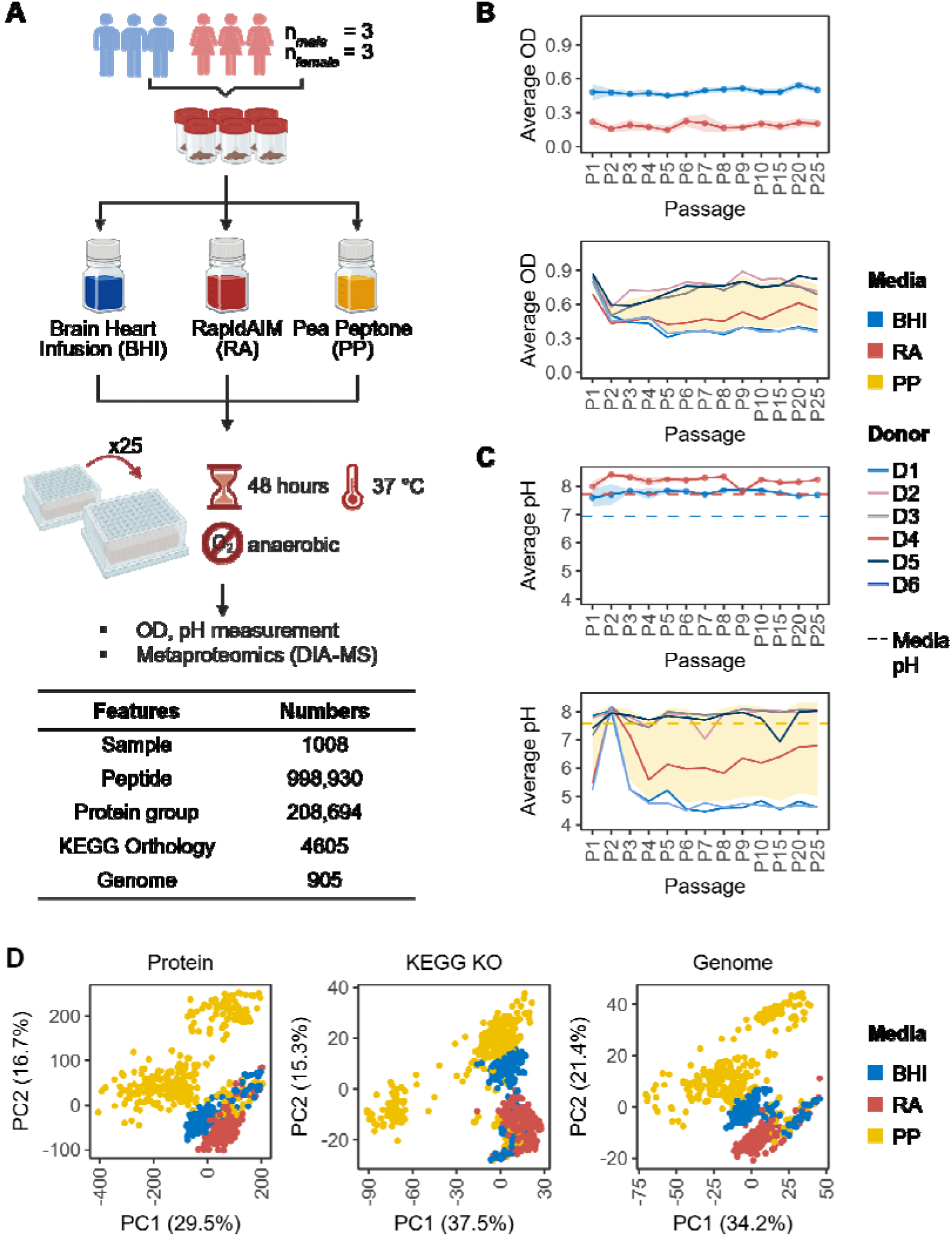
Experimental design and metaproteomic analyses of in vitro fecal microbial communities. (A) Experimental design. Six donor microbiomes were cultured and passaged every 48 hours for 25 passages in three media. Samples were collected at every passage for OD_600_ and pH measurements. Selected passages were analyzed with metaproteomics and a summary of metaproteomics results was indicated. (B) OD_600_ across passages. Top panel shows the BHI and RA groups, and ribbons represent the mean ± SD across all donors; bottom panel shows individual donor trajectories (colored by donor), which is overlaid with the overall mean ± SD ribbon across all donors. (C) pH across passages, plotted as in (B). Dashed lines indicate the pH of the respective medium as a reference. (D) PCA score plots of metaproteomic profiles at the protein, KO and genome levels.

## Results

### Fecal-derived *in vitro* microbiomes and metaproteomic analysis

In this study, six healthy adult donor microbiomes were cultured in three different media, including BHI and RapidAIM (RA) medium, as well as Pea peptone (PP) broth, an nutrient-rich non-animal medium that is commonly used for probiotic manufacturing [26]. Following a workflow similar to that was previously described [19], individual microbiomes were inoculated into each medium, cultured anaerobically in 96-deep-well plates, and passaged every 48 hours (Figure 1A). Microbial growth was monitored by measuring optical density (OD_600_) and pH at each passage. As shown in Figure 1B, microbiomes cultured in BHI and RA exhibited minimal inter-donor variation in OD_600_ across all passages, whereas those in PP showed substantially greater donor-dependent variability (Figure 1B). Overall, BHI (OD_600_ = 0.50 ± 0.04) and PP (0.60 ± 0.20) supported higher biomass yield than RA (0.19 ± 0.06), indicating that BHI and PP promote more rapid microbial growth. Similarly, consistent pH values were maintained in BHI (pH 7.69 ± 0.16) and RA (pH 8.12 ± 0.18) cultures across donors and passages, whereas PP cultures displayed marked donor-dependent variations (Figure 1C). Specifically, *in vitro* microbiomes derived from donors D1, D4, and D5 exhibited markedly lower pH values, along with lower OD_600_. Together, these results indicate that while PP elicits donor-specific variations in microbial growth and metabolic responses, BHI and RA support consistent *in vitro* fecal microbiome growth across all donors, suggesting the impacts of both donor microbiome composition and culture medium in the establishment of *in vitro* microbial communities with serial passaging.

To gain deeper insights into the community establishment, samples collected from passages P0–P10, P15, P20, and P25, were analyzed by metaproteomics using a DIA-MS workflow on the Orbitrap Astral MS system. In total, DIA-MS data were acquired from 1,008 microbiome samples, together with 9 quality control (QC) runs in which an identical QC sample was analyzed at different LC-MS batches throughout the study to monitor analytical reproducibility. Overall, 3.19 TB of DIA-MS data (1017 raw fles) were subjected for microbial peptide and protein identification and quantification, followed by taxonomic, functional annotations with MetaPilot, and taxon-function linked analyses with MetaX (see Methods). A total of 998,930 unique peptide sequences were identified, with a median of 60,973 peptides per sample (IQR: 43,043–88,693), resulting in the identification of 208,694 protein groups and 905 microbial genomes in the entire dataset (Figure 1A). Eleven out of the 1,017 DIA-MS datasets were excluded for the analysis because of insufficient data quality (<10,000 identified peptides or sample contaminations). We first assessed analytical QC samples using principal component analysis (PCA), which showed tight clustering of all QC samples (Figure S1A). Consistent taxonomic profiles across QC samples further confirmed high analytical consistency of the metaproteomics workflow in this study (Figure S1B).

To provide an overview of the metaproteomic data, we then performed PCA analyses for all samples at both protein-, function- and genome- levels. For functions, the Kyoto Encyclopedia of Genes and Genomes (KEGG) ortholog (KO) annotations of all proteins were used, resulting in 4,605 KOs quantified. These analyses demonstrated that BHI-and RA-derived *in vitro* microbiomes formed tight clusters, whereas PP-derived microbiomes showed markedly greater dispersion and shifted away from both BHI and RA groups (Figure 1D). This distinct and consistent clustering of samples according to culture medium suggests its profound impacts on the establishment of fecal microbial communities *in vitro* and reinforce the high metaproteomic data quality.

### Assessing donor and medium impacts on the *in vitro* microbiomes

To investigate the metaproteomic changes with microbiome passaging, we displayed the KO-level PCA score plot by passage, which showed that BHI communities gradually diverged from baseline (P0) to P3 and stable onward, whereas RA communities remained closer to P0 throughout passages (Figure 2A). A distinct sub-cluster emerged for PP communities starting P5, primarily comprising donors D1 and D5 (Figure 2A). Euclidean distances calculated based on the PCA score plots showed the least divergence of *in vitro* microbiomes cultured in RA to uncultured fecal inoculum (P0 microbiome) and the highest for those in PP (Figure 2B and Figure S2). Analysis of distances to previous passage further supported these observations, revealing small passage-to-passage changes in both BHI and RA microbiomes from P3. A dramatic shift was observed for microbiomes cultured in PP from P4 to P6, due to the sub- clustering of D1 and D5 (Figure 2B and Figure S2), which is consistent with the findings of distinct OD_600_ and pH patterns for both microbiomes in PP medium.

**Figure 2.**
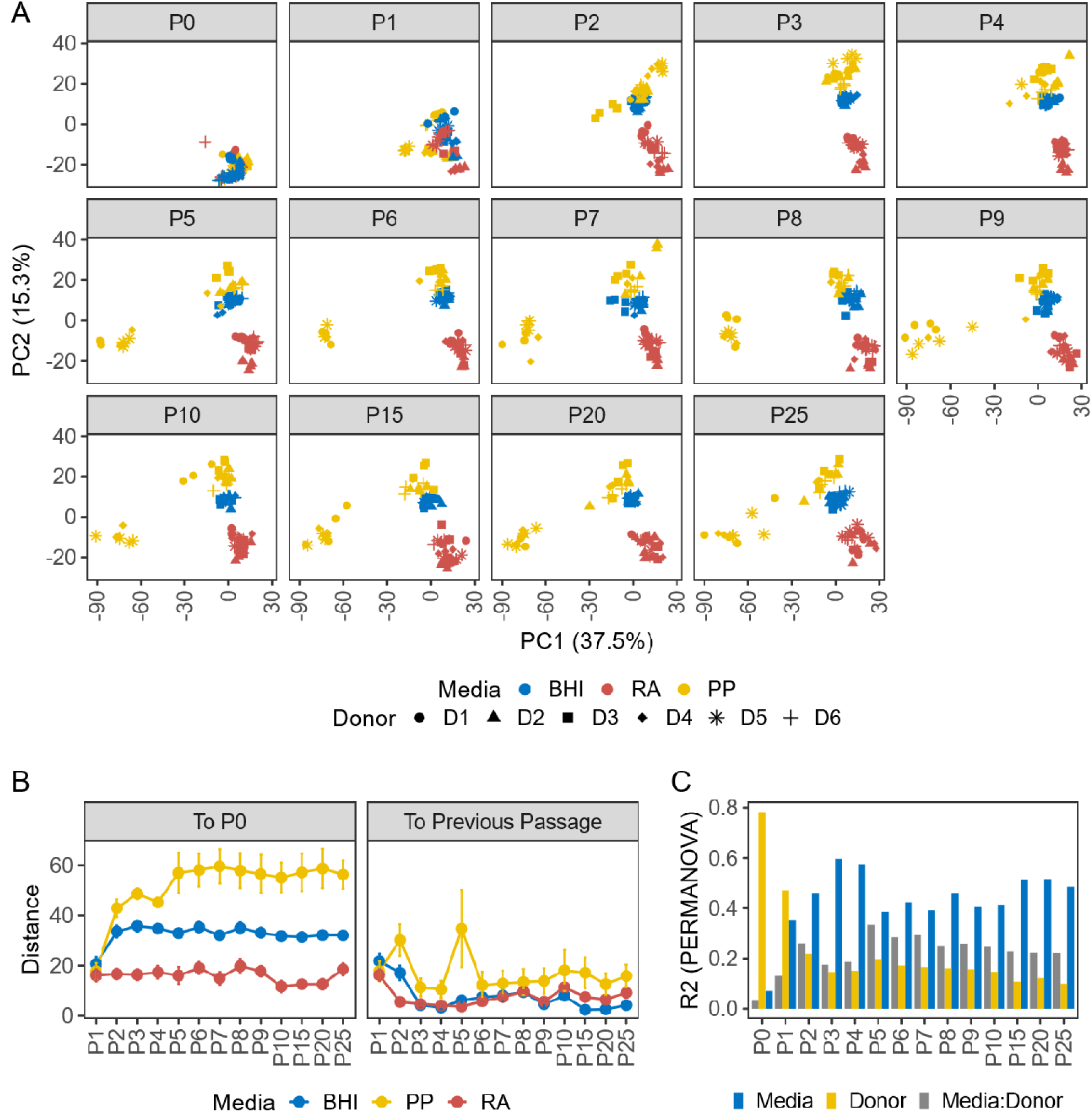
Metaproteomes of in vitro cultured microbiomes with different media. (A) PCA score plots based on KEGG KO profiles, faceted by passage. Colors indicate culture media, and symbols indicate donor microbiomes. (B) Distance analysis based on PCA score plot of KO profiles. Euclidean distance of each community to the corresponding P0 microbiome, and to the previous passage were shown in different panels. Colors indicate culture media. (C) PERMANOVA of KO profiles showing the proportion of variance explained (R²) by culture medium, donor, and their interaction at each passage. PCA components explaining 80% of the total variance were used for the analysis. Bars are colored according to the effect factor.

To statistically evaluate the contributions of medium and donor to the metaproteomic changes, we performed permutational multivariate analysis of variance (PERMANOVA). Across all samples, both culture medium and donor significantly influenced fecal-derived *in vitro* metaproteomes, with the culture medium explained a greater proportion of the variance than donor (KO level: R² = 0.19915, *p* < 0.001, pseudo-*F* = 351.5597 vs. R² = 0.13508, *p* < 0.001, pseudo-*F* = 95.3809; protein level: R² = 0.18886, *p* < 0.001, pseudo-*F* = 447.0187 vs. R² = 0.14860, *p* < 0.001, pseudo-*F* = 140.6921). A significant donor-medium interaction was also observed, indicating that individual donor microbiomes responded differently to culture conditions. PERMANOVA analysis conducted by passage showed passage-wise dynamics in the relative contributions of these factors (Figure 2C). Donor accounted for a greater proportion of the variance during the initial passages (P0 and P1), whereas from P2 onward, culture medium consistently explained more variance than either donor or the donor-medium interactions (Figure 2C). Notably, the impacts of both donor and culture medium remained statistically significant throughout all passages.

Similar findings were observed when protein-level abundance data was used for the PCA and PERMANOVA analyses (Figure S3-5). Collectively, these results demonstrate that both the initial donor microbiome and culture medium shape the functional trajectories of *in vitro* microbial communities, with the culture medium being the primary factor driving the variations of metaproteomic composition. Briefly, RA most effectively maintained the functional profiles of original donor microbiomes, while significant changes were observed in microbiomes cultured in PP medium.

### Metaproteome-based taxonomic profiling of *in vitro* cultured microbiomes

Metaproteomics provides direct biomass-based taxonomic characterization of microbial communities. In this study, the MGnify human gut microbial genome catalog database was used for gut microbial peptide identification followed by an adaptive genome selection with MetaPilot [24]. In total, 905 species-representative genomes from 10 phyla, 15 classes, 40 orders, 86 families, and 317 genera were identified in this study. Taxonomic profiling across all donors and by individual donor microbiomes revealed distinct, culture medium-dependent community structures, while still visible donor-specific differences (Figure 3A, Figure S6 and S7). Among the three media, RA maintained a higher relative abundance of Bacteroidetes, including the *Bacteroidaceae* family, compared to both BHI and PP. BHI was characterized by a profound enrichment of Proteobacteria, particularly members of the *Enterobacteriaceae* family, consistent with previous findings that nutrient-rich BHI medium preferentially supports the rapid expansion of fast-growing facultative anaerobes [19]. PP also promoted the growth of Proteobacteria, but exhibited a distinct donor-dependent pattern in which Proteobacteria increased and peaked around P3-P4 and subsequently declined in D1 and D5, which was accompanied by a substantial expansion of Firmicutes, particularly the *Enterococcaceae* family (Figure S6 and S7).

**Figure 3.**
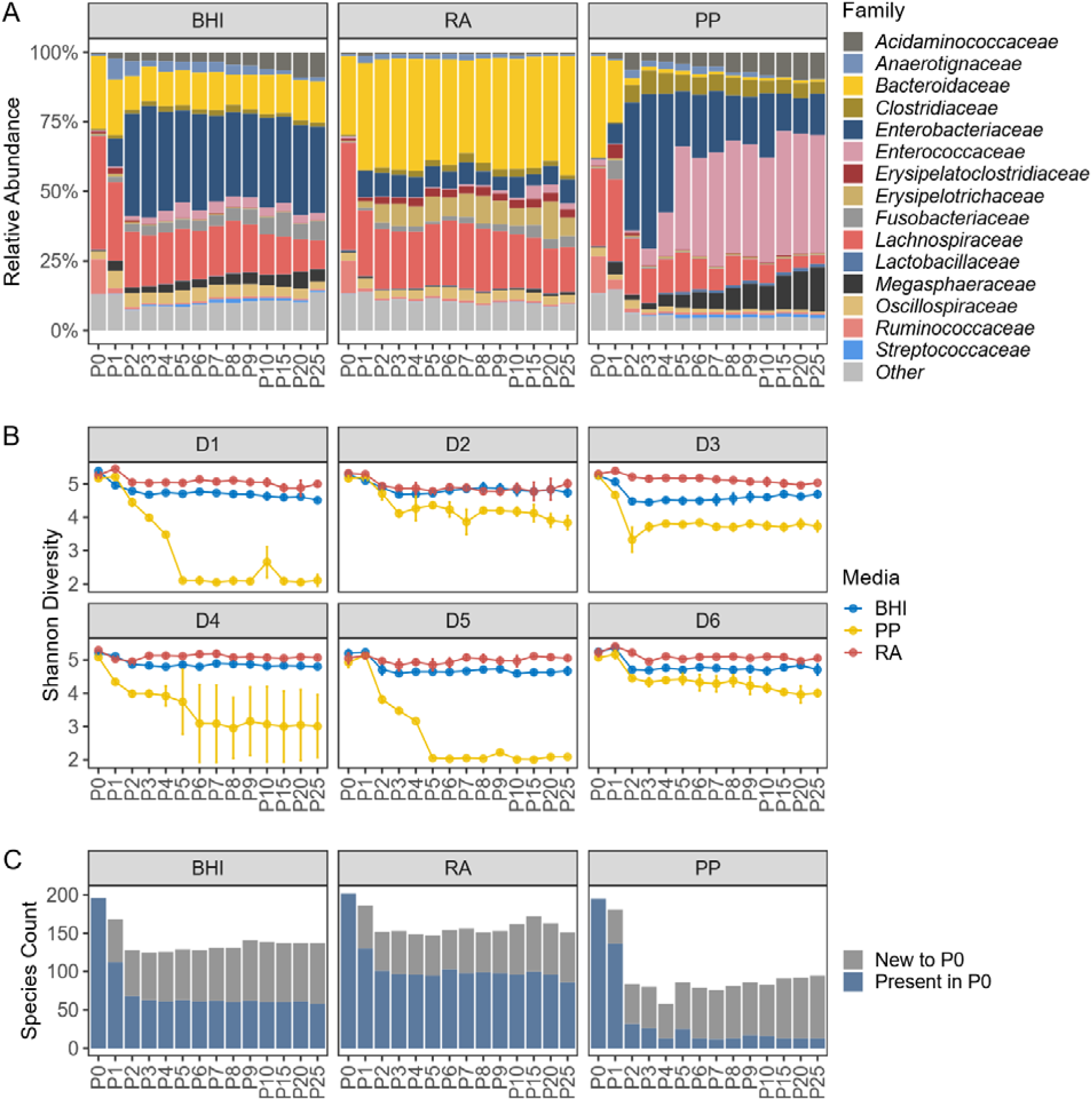
Taxonomic composition and diversity of in vitro microbiomes. (A) Family level composition of group means at each passage, faceted by culture media. (B) Shannon diversity calculated with species abundance in each donor and passage. Colors indicate culture media, and mean ± SD of culture replicates was shown. (C) Species retention analysis across all donors. Stacked bars show species retained from P0 (dark blue) and species not detected at P0 microbiomes (grey).

Shannon diversity index was calculated at the species level across all passages, which showed that the microbiomes cultured in BHI and RA retained stable species diversity over time with only slight reduction with passaging (Figure 3B). In contrast, PP displayed greater donor-dependent variability, with donors D1 and D5 showing marked decreases in diversity, which could largely attribute to the increased dominance of *Enterococcus faecalis* in these *in vitro* microbiomes in later passages.

We next evaluated the extent and which microbial species were retained during serial passaging. In this analysis, we applied a three-level filtering strategy to ensure that only confidently identified species were considered (see Methods). The results showed that the total number of species remained comparable to that of the baseline (P0) microbiomes in both BHI and RA groups, whereas a marked reduction was observed in the PP group (Figure S8). The evaluation of species identity presents in P0 and later passages showed that, in all donors, RA retained the largest proportion of P0 species, whereas PP retained the fewest (Figure S8). Assessment of the overall species retention across all the six donors (a species was considered as present in a passage if it was identified in at least 3 donors) again showed that RA preserves the greatest proportion of P0 species (57%- 66.9% species in P2 to P25) followed by BHI (42.3%- 53.1% species in P2 to P25) (Figure 3C).

Altogether, the functional and taxonomic profiling analyses showed that the microbiomes underwent marked donor-dependent shifting during serial passaging in PP medium. In contrast, microbiomes cultured in BHI and RA media maintained stable taxonomic compositions in later passages while preserving community diversity.

### Metaproteomic evaluation of community dynamics of *in vitro* microbiomes

In the serial passaging experiment, four replicates per donor and medium were cultured and passaged to the corresponding replicates in the following passage, generating four culture lineages for each condition. To investigate the community stability, we compared the variations within lineage across successive passages with those between lineages in the same passage. We defined stable passaging or stable *in vitro* community (SIC) as a condition in which the within-lineage dissimilarity was not significantly greater than the inter-lineage dissimilarity of that passage. Two different dissimilarity matrices, Bray-Curtis and UniFrac, were calculated with species composition quantified with metaproteomics. For this analysis, samples in the PP group were removed due to the marked community shifting and diversity loss as demonstrated above. As shown in Figure 4A, BHI communities showed no significant difference from P5 for both Bray-Curtis and UniFrac analyses, while RA achieved no significance starting from P4, and as early as P3 for weighted UniFrac analysis. Overall, the within-lineage dissimilarities across successive passages remained low from P5 for both BHI and RA conditions. Although a slight increase of the within-lineage dissimilarities was observed at later passages, they remained not significantly greater than the corresponding inter-lineage dissimilarities, suggesting that fecal-derived SICs can be achieved using both BHI and RA media.

**Figure 4.**
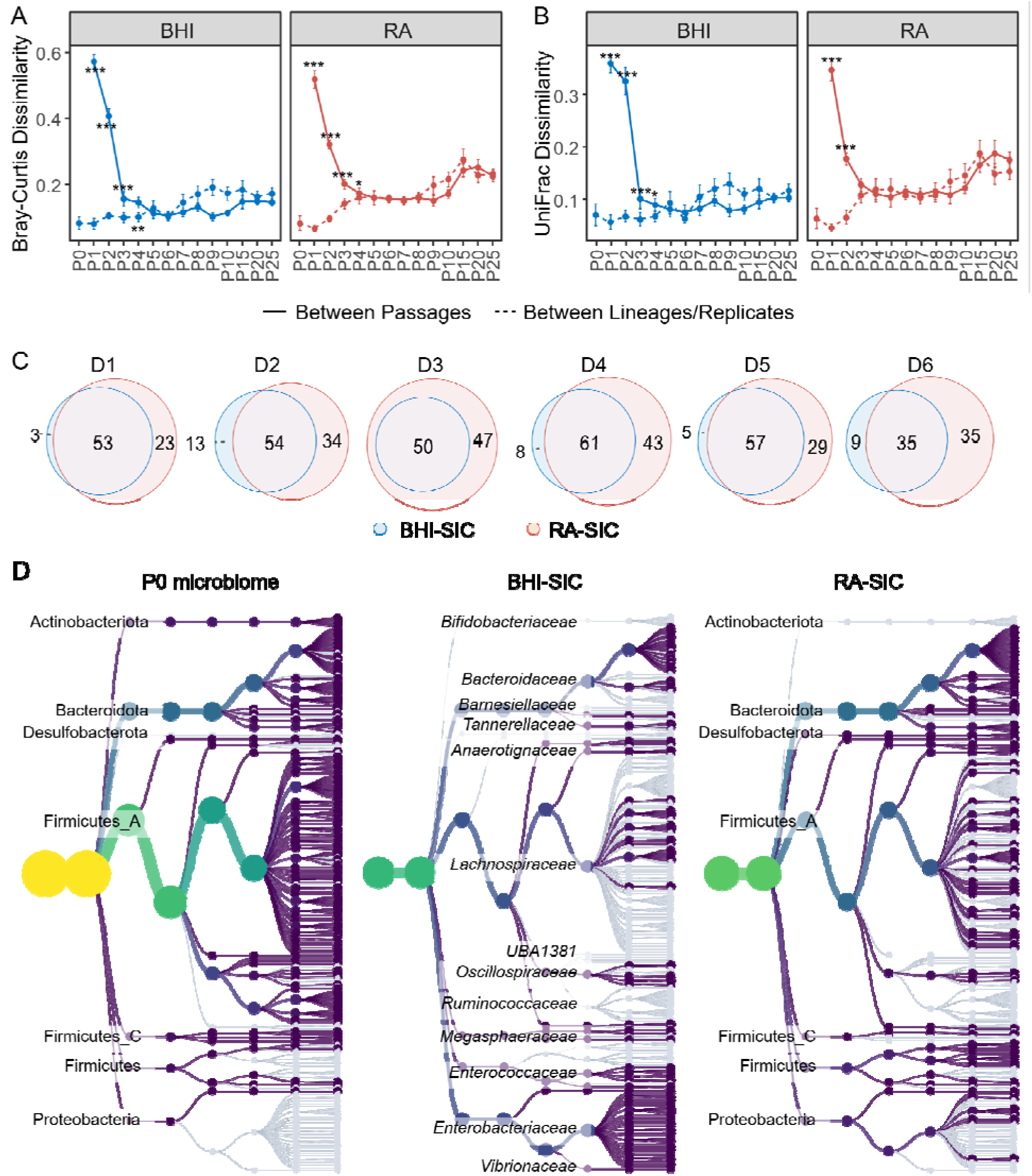
Stability and species retention in in vitro microbial communities. (A) Within- and inter-lineage Bray-Curtis dissimilarities of species compositions. Error bar shows the standard deviation of the six donors. ***, BH-adjusted p <0.001 with one-sided Wilcoxon test. (B) Within- and inter-lineage UniFrac dissimilarities of species compositions. ***, BH-adjusted p <0.001 and **, BH-adjusted p <0.01 with one-sided Wilcoxon test. (C) Venn diagrams showing the overlap of retained P0 species in BHI- and RA-SICs. (D) Phylogenetic tree showing the presence (colored) and absence (grey) of confidently identified species in P0, BHI-SICs and RA-SICs, respectively. The same tree was used for all three groups, and phylum or family names were indicated on the tree plots when possible.

To further assess which P0 microbiome species were retained in SICs, microbiomes from P5 to P7 of both RA and BHI groups were selected as representative SICs, thereafter referred to as RA-SIC and BHI-SIC, respectively. For confident species identification, the same three-level filtering strategy was used and a species was considered retained in the SICs if it was detected in at least two of the three representative passages. Figure 4B showed the overlap between retained P0 species in BHI- and RA-SICs in each donor microbiome, suggesting that RA consistently retained a greater number of P0 species across all donors. Notably, most species retained in BHI-SICs were also retained in RA-SICs. Overall, 158 of the 207 P0 species persisted in the stable SICs, including 94 shared between BHI- and RA-SICs, with 52 species uniquely retained in RA-SICs. Phylogenetic display of the P0, BHI-SIC and RA-SIC species showed that *Enterococcaceae* species were low in P0 microbiomes, but was enriched in BHI-SICs, while *Bifidobacteriaceae* species were lost in both BHI- and RA- SICs (Figure 4C). Compared to BHI-SICs, RA-SICs retained more species belonging to the *Lachnospiraceae* and *Bacteroidaceae* species (Figure 4C), consistent with the observations in taxonomic composition.

### Taxon-specific functions driving differential SIC establishments

To identify the functions that drive the differences between BHI- and RA-SICs, we applied a partial least squares discriminant analysis (PLS-DA) using KO abundance data. PLS-DA analysis showed significant differences between BHI- and RA-SICs with a model goodness of prediction (Q^2^) value of 0.94. A total of 1069 differentially abundant KOs were identified using a variable importance in projection (VIP) value threshold of 1 (Table S1). Figure 5 showed the heatmap of top 50 VIP KOs in both the BHI-SIC, RA- SIC, and the P0 microbiomes as a reference for uncultured fecal inoculum. Group clustering of these top 50 VIP KOs showed that RA-derived SICs clustered most closely to P0, consistent with the PCA analysis. There are two KO clusters with cluster 1 having higher abundances in BHI-SIC, while cluster 2 having higher abundances in P0 and RA- SIC (Figure 5).

**Figure 5.**
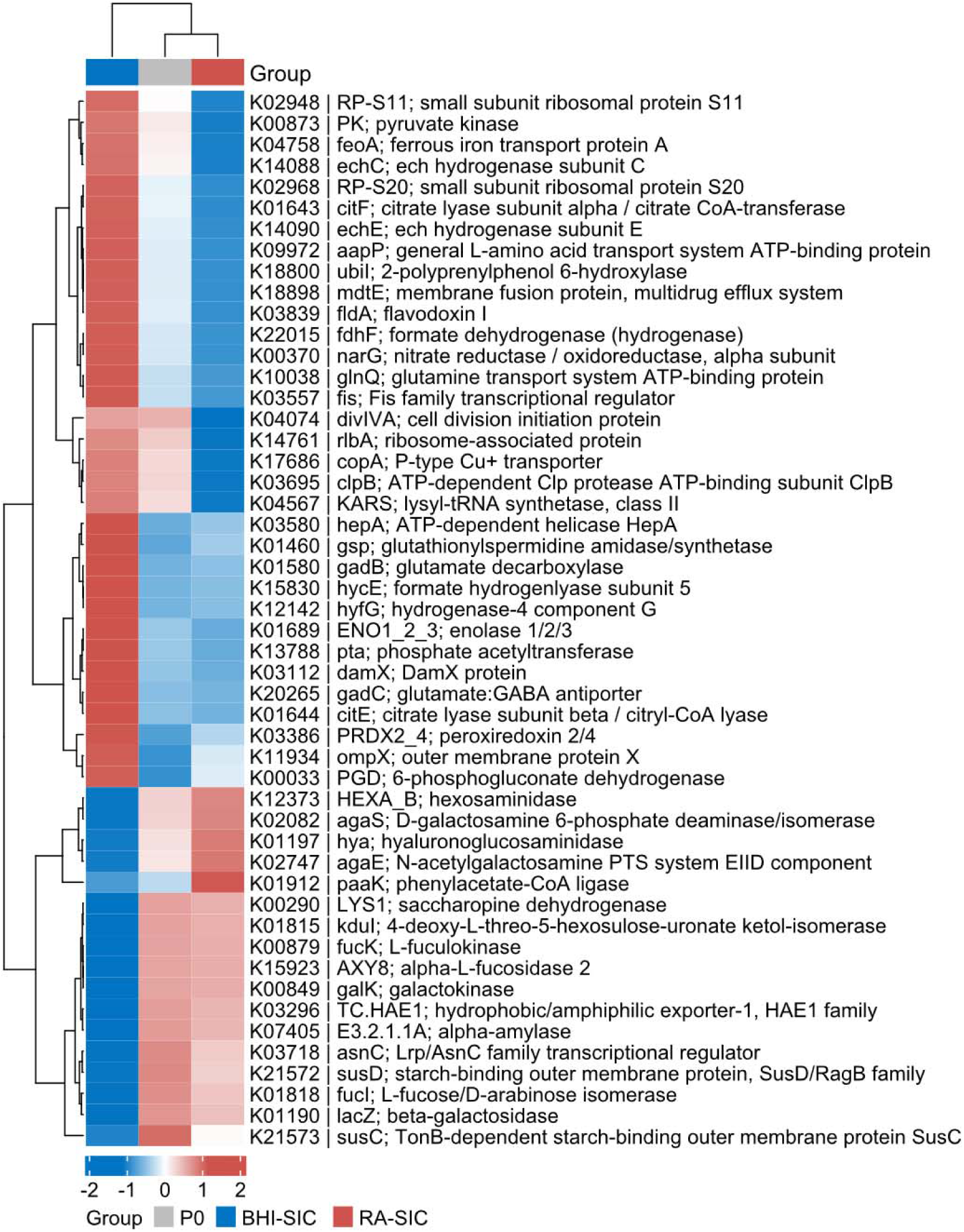
Heatmap of top differentially abundant KOs between BHI- and RA-SICs. The KOs were selected based on PLS-DA VIP. Values represent group means in the heatmap, and P0 group was included as reference. Both rows and columns were clustered using Euclidean distance. The K number, gene symbol, and name are shown for each KO.

The up-regulated KOs in BHI-SIC (cluster 1) are mainly associated with microbial stress adaptation mechanisms. These include Flavodoxin I (K03839), peroxiredoxin 2/4 (K03386), nitrate reductase (K00370), glutamate decarboxylase (K01580), and ATP-dependent Clp protease ATP-binding subunit ClpB (K03695), which are characteristics of microbial responses to acid, oxidative stress and nutrient limitation [27–29]. On the contrary, KOs upregulated in RA-SIC and P0 (cluster 2) were enriched in pathways associated with carbohydrate metabolism, in particular the metabolism of mucin glycans, such as galactokinase (K00849), hexosaminidase (K12373), L-fuculokinase (K00879), L-fucose/D-arabinose isomerase (K01818), alpha-L-fucosidase 2 (K15923), D-galactosamine 6-phosphate deaminase (K02082), N-acetylgalactosamine PTS system EIID component (K02747), and starch-binding outer membrane proteins (K21573, K21572). These findings are consistent with the distinct nutrient environments, as RA medium and the *in vivo* gut environment are enriched in mucin as a major nutrient source [10, 30], while BHI is rich in non-specific animal derived amino acids.

One advantage of metaproteomics is the direct link of taxa to expressed functions in microbial communities. Using an operational taxon-function (OTF) analysis framework in MetaX [31], we linked 3736 KOs to 203 genera, yielding 45,664 genus-KO OTFs in this study. PLS-DA analysis identified 853 OTFs that are up-regulated and 634 down- regulated in RA-SICs compared to BHI-SICs. By matching the top 50 VIP KOs in Figure 5, we found that those most significantly enriched functions in RA-SICs are mainly attributed to the mucin-utilizing species *Bacteroides* and *Phocaecola* [32], while those most significantly enriched in BHI-SICs are mainly attributed to *Escherichia* (Figure S9). These taxon-specific functional changes align with the observed taxonomic profiles, and highlight that the functions associated with complex glycan metabolism and stress response may drive the differential SIC establishment in BHI and RA media.

The increase of complex glycan metabolism enzymes in RA-SICs is consistent with the well-recognized role of the gut microbiota in fermenting complex indigestible carbohydrates, including dietary fibers and mucins. To further characterize carbohydrate metabolism in RA-SICs, the identified microbial proteins were annotated with the CAZyme database, resulting in the assignment of 3713 proteins to 76 CAZyme families. As shown in Figure 6A, the uncultured P0 microbiomes contain the highest percentage of CAZyme protein abundance (median: 1.57%, IQR: 1.33% – 1.77%), which is followed by RA-SICs (median: 1.22%, IQR: 1.22% – 1.37%). The BHI-SICs exhibited the lowest CAZyme abundance (median: 0.40%, IQR: 0.33% – 0.46%), consistent with the nutrient-rich composition of BHI medium, which contains few complex carbohydrates. Across all samples, GT35, GH20, and CBM48 were the most abundant Glycosyltransferase (GT), Glycoside Hydrolase (GH), and Carbohydrate-Binding Module (CBM) families, respectively (Figure 6B). Notably, although the overall abundance of CAZyme proteins was lower in RA-SICs than in P0, GH20 was substantially enriched in RA-SICs (Figure 6B). Members of the GH20 family primarily include mucin glycan-degrading enzymes, such as β-*N*-acetylhexosaminidases and β- *N*-acetylgalactosaminidases, consistent with the enrichment of mucin degradation functions in RA-SICs as demonstrated above.

**Figure 6.**
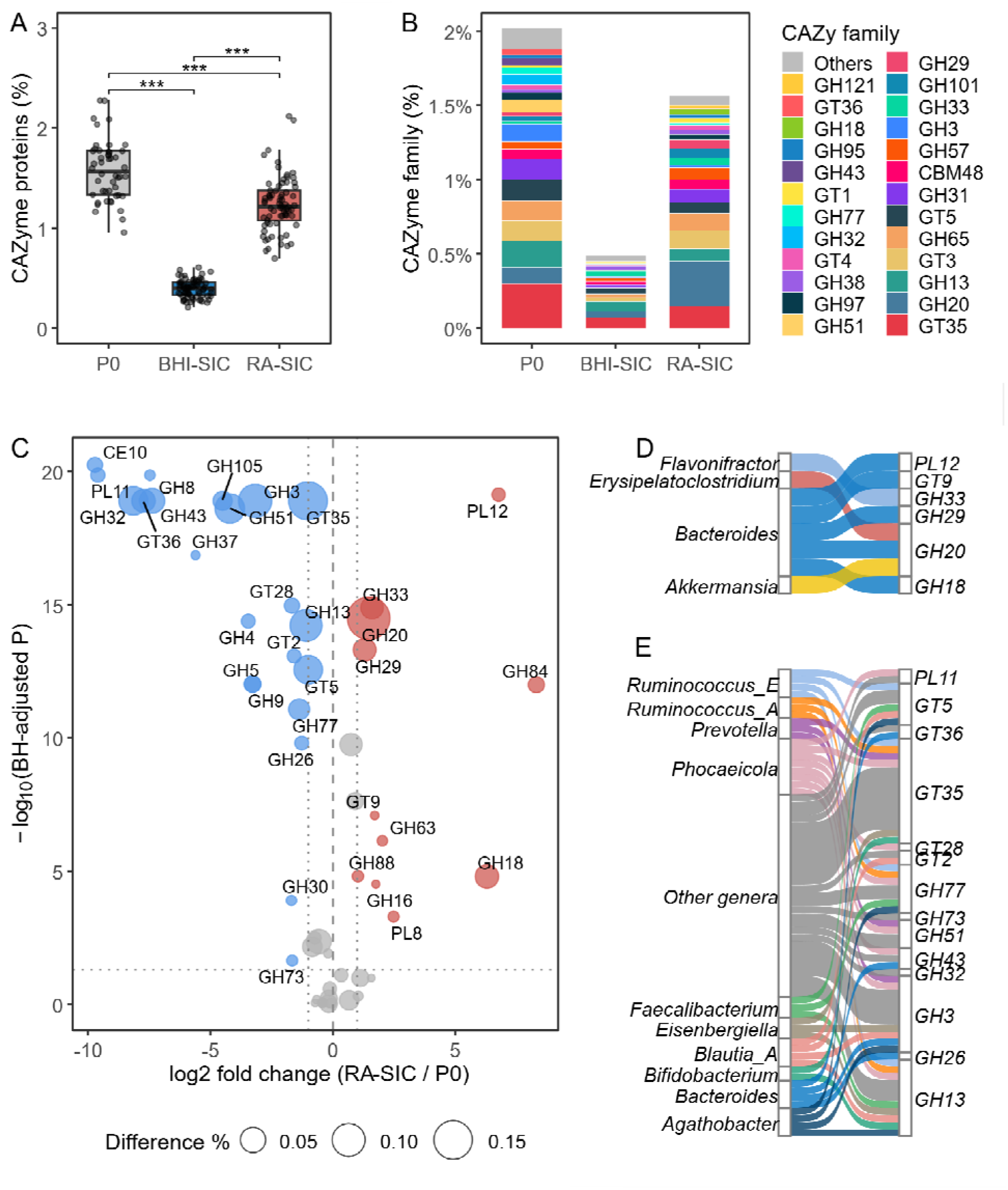
CAZyme expressions in fecal-derived SICs and P0 microbiomes. (A) Proportion of total CAZyme proteins in metaproteomes of P0 microbiome and fecal-derived SICs. ***, BH-adjusted p <0.001 with pairwise Wilcoxon test. (B) Relative abundance of CAZyme families in P0 microbiome and fecal-derived SICs. (C) Volcano plot showing the differentially abundant CAZyme families between P0 and RA-SICs. Size of the bubbles show the absolute differences between two groups. CAZyme families with BH-adjusted P < 0.05 and FC > 2 were indicated in the plot (red as up- regulated in RA-SICs and blue as up-regulated in P0. (D-E) Significantly different CAZyme-genus OTFs that matched to the identified differential abundant CAZyme families in Figure 6C. Ribbons are colored based on taxa and size corresponds to the number of significant OTFs.

Differential abundance analysis identified 11 CAZyme families that were significantly enriched and 22 that were depleted in RA-SICs compared to P0 (Benjamini-Hochberg (BH)-adjusted p < 0.05 and fold change (FC) ≥ 2; Figure 6C). In addition to GH20, the families GH33 (sialidases and trans-sialidases hydrolyse), GH29 (α-fucosidases), GH18 (chitinases and β-N-acetylglucosaminidases), GH84 (β-N-acetylglucosaminidases and β-N-acetylhyaluronidases), and PL12 (heparin or heparin-sulfate lyase) were also significantly enriched in RA-SICs. CAZyme-genus OTF analysis showed that the enriched CAZyme families in RA-SICs, in particular the mucin glycan metabolism enzymes, were primarily contributed by *Bacteroides, Erysipelatoclostridium, Akkermansia,* and *Flavonifractor* (Figure 6D). The CAZymes enriched in uncultured P0 microbiomes were attributed to a wide range of microbial genera, including common commensal gut microbes (Figure 6E), which is consistent with the diverse nature of human gut nutrient environment. Altogether, these functional characterizations suggest that complex carbohydrate utilization is a crucial factor and the presence of mucin glycan allows to preserve key functionality and compositions of the native gut microbiome. In contrast, nutrient-rich conditions that support rapid microbial growth may promote a transition toward a stress-adapted functional state and lead to the dominance of fast-grower in the community [20].

### Metaproteomic measurements of microbial risk factors in SICs

Like shotgun metagenomics, metaproteomics does not target specific marker proteins, enabling the potential to detect any expressed or secreted proteins present in a sample, provided that their sequences are included in the database and the identification false discovery rate (FDR) is well controlled [21, 33–35]. In this study, we leveraged this advantage to profile microbial risk factors that are commonly considered in the safety assessment of biotherapeutics and probiotics, including virulence factors (VF), antimicrobial resistance proteins (ARP), and bacteriophage proteins. Each of the databases was concatenated with either the identified microbial peptide database or the complete UHGG database to perform targeted database searches to identify microbial risk factor proteins (see Methods). Comparison of the relative abundance of risk factor proteins within the metaproteomes revealed that both BHI- and RA-SICs displayed significantly higher proportions of VFs and ARPs than the corresponding P0 microbiomes, with the greatest increase observed in BHI-SICs. In contrast, both SIC models exhibited lower proportions of bacteriophage proteins than the P0 microbiomes (Figure 7A)

**Figure 7.**
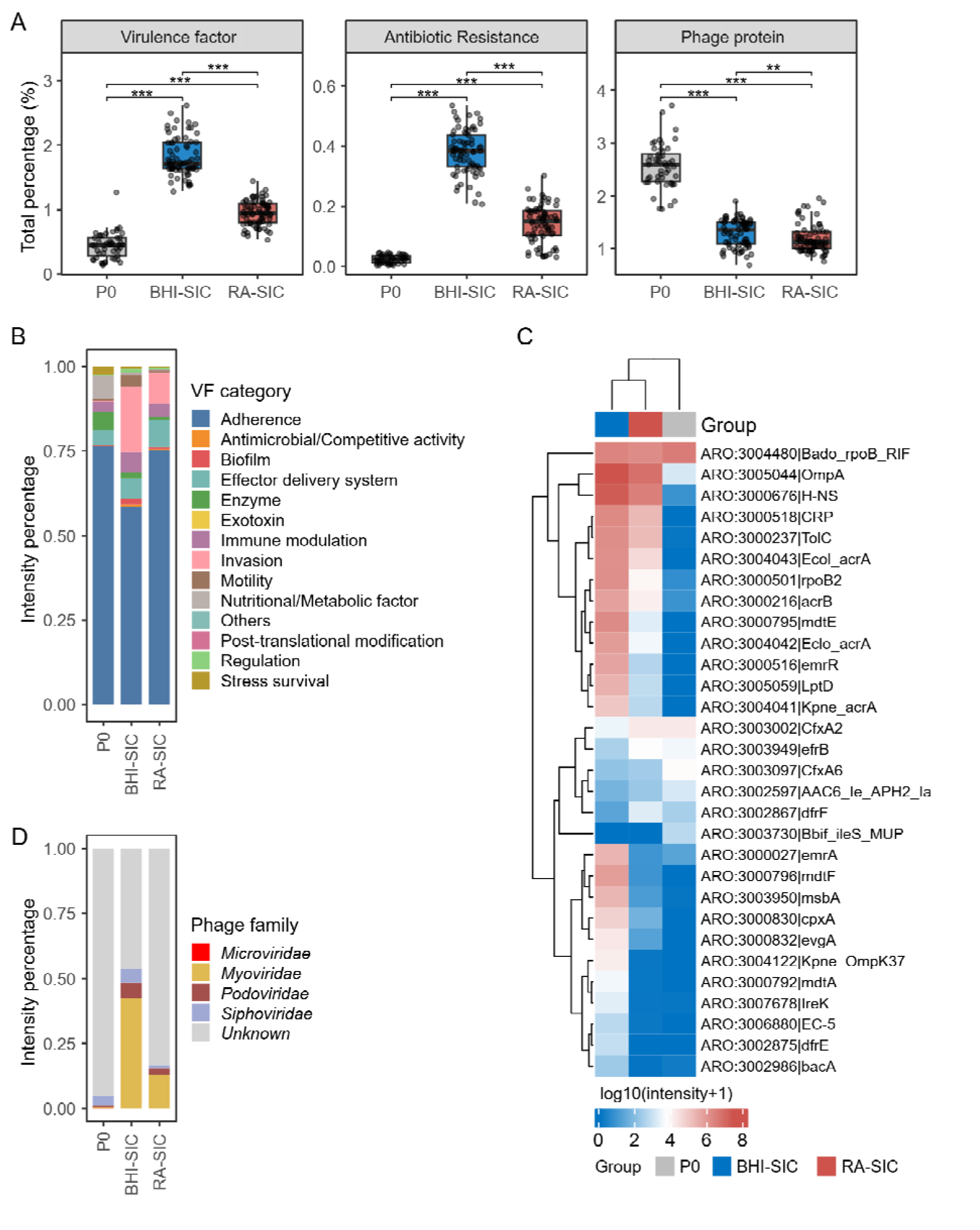
Microbial risk factors in fecal-derived SICs and P0 microbiomes. (A) Total relative abundance of each microbial risk factor type of proteins in total metaproteomes. MS data were searched against databases containing the Gut Phage Database (GPD), Virulence Factor Database (VFDB), and antibiotic-resistance protein database (ARP). The sum protein intensity of all identified risk factor proteins was used for calculation of relative abundance in total metaproteomes. ***, BH-adjusted p <0.001, **, BH-adjusted p<0.01 with pairwise Wilcoxon test. (B) Mean group intensity of virulence factor composition by VF category. (C) Heatmap of the abundances ARP proteins detected in ≥50% of samples in at least one group. Values represent group means in the heatmap. Both rows and columns were clustered using Euclidean distance. (D) Family level composition of identified phage proteins.

A total of 516 VF proteins were identified across all samples. We showed that proteins involved in adherence constituted the most abundant VF category in both P0 microbiomes and SICs. However, proteins associated with invasion and biofilm were markedly enriched in both BHI- and RA-SICs, particularly in BHI-SICs. BHI-SICs also exhibited a pronounced increase in VFs associated with motility. We also found that the proteins involved in enzyme and nutritional/metabolic factors were reduced in both SICs compared to those of P0 microbiomes (Figure 7B).

This study identified 89 ARPs of which 30 were detected in at least half of the P0 microbiome, BHI-SIC, or RA-SIC samples (Figure 7C). Bado_rpoB_RIF, which confers resistance of commensals, such as *Bifidobacterium adolescentis*, to rifampin, is among the most abundant ARPs in all the samples regardless of culture conditions. Several ARPs displayed consistent trends of changes in both SICs, including the increase of antibiotic efflux/influx regulation ARPs of Gram-negative bacteria, such as OmpA, H-NS, CRP and TolC, and the decrease of antibiotic inactivation ARPs, such as CfxA6. Nevertheless, majority of ARPs remained at a comparable abundance level between the P0 microbiomes and RA-SICs (Figure 7C), indicating that the overall antimicrobial resistance profile was largely preserved in RA-SICs.

A total of 1619 phage proteins were identified in the current dataset. Consistent with the gut phage database used for phage protein identification [36], most identified phage proteins could not be assigned to a known taxonomic group (Figure 7D). Majority of the annotated phage proteins belonged to the *Caudovirales* order, including *Myoviridae*, *Siphoviridae*, and *Podoviridae*, which is in agreement with previous findings [37]. Compared with the P0 microbiomes and RA-SICs, BHI-SICs exhibited a higher relative abundance of *Myoviridae* proteins and a lower proportion of unclassified phage proteins (Figure 7D). Previous studies have reported that *Myoviridae* possesses a broader host range with majority of them are Firmicutes and Proteobacteria [36, 37]. This is consistent with the observations that BHI-SICs were characterized by a marked enrichment of Proteobacteria and less representation of Bacteroidota. Altogether, RA- SICs maintained a profile of microbial risk factors that more closely resembled the P0 microbiomes when compared to those of BHI-SICs.

## Discussion

The nutrient composition of culture medium is crucial to maintain the *ex vivo* microbiomes and a key factor for the establishment of *in vitro* microbial communities [20, 38]. In this study, using a large-scale metaproteomic approach, we profiled the taxon-specific functional dynamics of human fecal microbiomes cultured in three culture media over 25 serial passages. We found that the medium choice strongly influenced the establishment, composition, functional profiles, and stability of *in vitro* fecal-derived microbial communities. SICs could be established using BHI and RA media, both of which have previously been shown to support *ex vivo* gut microbiome growth [19, 39], but marked differences in both taxonomic and functional compositions were observed. In contrast, *in vitro* serial culturing in PP medium, a food-grade medium developed to support beneficial probiotic organisms [26], resulted in substantial changes of the communities in a donor-dependent manner. Overall, these findings support the feasibility of generating fecal-derived SICs as microbiome models, while highlighting the critical importance of carefully selecting the culture medium to preserve desired microbiome characteristics.

Consistent with previous observations by Aranda-Diaz et al. [19], we found that the fecal-derived SICs with BHI medium displayed higher level of Proteobacteria, in paticular *Enterobacteroiacea* species. In contrast, RA-SICs displayed a much lower level of Proteobacteria, while having higher abundances of *Bacteroidaceae* species. These contributed to the higher species diversity, more baseline species retention, and higher similarity of RA-SICs to the uncultured microbiomes, compared to BHI-SICs. BHI is a nutrient-rich medium that supports the growth of a broad range of microorganisms, including fastidious or pathogenic species. This is consistent with the expansion of *Enterobacteriaceae*, including *E. coli*, in BHI-SICs, which may rapidly consume available nutrients and thereby gain a competitive advantage over other commensal microbes in the microbiome [20]. RA differs substantially in nutrient composition to BHI, with mucin providing complex glycans that more closely resemble the nutritional environment of the gut [11, 39]. The utilization of mucin glycans requires a spectrum of enzymes from different microbial species, providing one mechanism underlying the complex microbe-microbe and host-microbe interactions of the intestinal ecosystem [32, 40]. Accordingly, metaproteomic analysis in this study demonstrated that, compared with BHI-SICs, proteins most significantly upregulated in RA-SICs were enzymes involved in the degradation and metabolism of mucin O-glycan components, including galactose, fucose, N-acetylgalactosamine, galactosamine, and sialic acid. Taxon-specific functional analysis further revealed that mucin-utilizing species *Bacteroides* and *Phocaecola* largely contributed to the up-regulation of these proteins, consistent with their increased abundances in RA-SICs and less representation in BHI-SICs. CAZyme focused analysis further supported these findings, showing that BHI-SICs had the lowest overall CAZyme levels, whereas RA-SICs more closely resembled the uncultured microbiomes. Additionally, we also found that the uncultured microbiomes displayed higher level and broader families of CAZymes, while RA-SICs were characterized by the enrichment of mucin glycan metabolism related CAZymes. This observation suggests that glycans beyond mucin, including dietary fibers and other host-derived complex carbohydrates [41, 42], may also contribute to maintaining the functional and compositional complexity of the native microbiome in human gut.

Both BHI and PP are nutrient-rich media, but with animal- and plant-based nutrients, respectively. Consistent with their nutrient-rich nature, BHI- and PP-cultured microbiomes yielded higher biomass and relatively lower pH than those cultured in RA in all passages. Taxon-specific functional profiling showed that *Escherichia* gained a competitive advantage in BHI and early passages of PP through the activation of stress-response systems, including acid resistance, membrane reinforcement, and protein-folding and repair mechanisms. The stresses may result from rapid nutrient depletion by fast-growing microorganisms, leading to the accumulation of metabolic by-products that promote acidic and oxidative conditions in the cultures. In later PP passages, some donor microbiomes (e.g., D1 and D5) underwent a dramatic shift in community composition, with absolute dominance by *Enterococcus* species accompanied by a marked decline in pH to approximately 5 (Figure S10A). OTF analysis comparing early passages (P0–P2) with later passages (P4–P5) when *Enterococcus* exerted increasing abundances and replace *Escherichia* as dominant species showed that, instead of glutamate decarboxylation (GAD)-mediated acute acid resistance pathway adopted by *Escherichia*, *Enterococcus* utilized arginine deiminase (ADI) pathway (e.g., up- regulation of ADI and ornithine carbamoyltransferase (OTC)) for acid resistance (Figure S10B-C). Compared with the GAD pathway, which consumes intracellular protons by converting glutamate to GABA, the ADI pathway neutralizes protons through releasing ammonia and generates one ATP for bacterial growth [27]. The latter is more energy-efficient and may be better suited for sustainable growth under acidic conditions, potentially conferring long-term growth advantage of *Enterococcus*.

In addition to taxonomic composition and protein expressions, metaproteomics has an advantage to capture a broad range of proteins or proteoforms without targeting specific organisms or protein types [22, 34]. As long as the reference protein database is available and a suitable false discovery control strategy is applied, respective proteins can be identified and quantified. This advantage is particularly relevant to DIA-MS, in which theoretically all ions are analyzed and fragmented, enabling comprehensive detection of proteins, including low-abundance or less-characterized components [43, 44]. In this study, we demonstrated this capability by analyzing commonly considered microbial risk factors in biotherapeutic or probiotic assessments, including virulence factor (VF), antimicrobial resistant proteins (ARPs) and bacteriophage proteins in the SICs. We confidently identified VF, ARP and phage proteins in the current dataset and showed that the SICs displayed increased overall protein abundances of VF and ARP, and lower phage protein abundances, compared to native uncultured microbiomes. The increased VF and ARP in SICs may be due to the enrichment of Proteobacteria, which are generally associated with higher levels of these risk factors, or because the microbial risk factors are better characterized in cultivable bacterial species [45–47]. Compared to BHI-SICs, the RA-SICs showed lower levels of microbial risk factor proteins, consistent with their higher similarity to the native microbiome. Most bacteriophage proteins were assigned to unknown taxa without predicted hosts, limiting our ability to interpret their roles in community establishment and highlighting an important knowledge gap warrants future investigations.

Overall this metaproteomic study provides mechanistic understanding of the establishment of *in vitro* microbial communities from human fecal microbiomes, and reinforce the crucial role of culture medium in the community assembly. The results may inform further optimization of the culture medium to ultimately generate stable, scalable and diverse microbial communities that better recapitulate the native microbiomes for microbiome studies and translational applications. The current study is limited by the small number of donor microbiomes, and a larger cohort of both healthy and patient microbiomes may be needed to better define how donor microbiomes shape the *in vitro* community establishment. Additionally, metaproteomics is a rapidly evolving technology with current limitations in dealing with very large databases, in particular for DIA-MS data [34, 43]. To address this challenge, this study used a species-representative genome database and a multi-step database search strategy, however this approach may fail to capture protein variants, mutations, and strain-level differences that are common in human gut microbiomes [48]. Therefore, the acquired MS data may not be fully leveraged to capture the full breadth of microbes and expressed microbial proteins. As improved database search strategies and computational resources become available, the present dataset can be readily reanalyzed in a more mutation- and strain-aware framework to provide a more comprehensive understanding of the community dynamics.

## Methods

### Human stool collection and biobanking

Fecal microbiomes were obtained from three male (D1, D4, D6) and three female (D2, D3, D5) healthy donors (27 - 33 years old) and biobanked as previously described [10, 49]. Briefly, stool samples were collected in a deoxygenated glycerol buffer, kept on ice, and shipped to the laboratory for processing; immediately after receiving, the stool samples were transferred to anaerobic workstation, thoroughly homogenized, and top up with glycerol buffer to make 20% fecal slurry. The fecal slurry was then filtered to remove large particles and stored at −80°C until use. The stool collection protocols (#20160585-01 H and H-09-22-8466) were approved by the Ottawa Health Science Network Research Ethics Board at the Ottawa Hospital and the University of Ottawa Office of Research Ethics and Integrity, Ontario, Canada, respectively.

### Culture medium preparation

BHI was prepared by dissolving dehydrated powder (BD Bacto™) in double distilled water (ddH_2_O) to a final concentration of 37g/L according to the manufacturer’s instructions. The medium was heated to ensure complete dissolution and sterilized by autoclaving at 121 °C for 15 minutes.

RA was prepared following the protocol described by Li et al.[10] Briefly, the RA base medium is consisted of peptone water (2.0 g/L), yeast extract (2.0 g/L), sodium chloride (1.0 g/L), dipotassium hydrogen phosphate (0.4 g/L;), potassium dihydrogen phosphate (0.4 g/L), magnesium sulfate heptahydrate (0.1 g/L), calcium chloride dihydrate (0.1 g/L), sodium bicarbonate (4.0 g/L), sodium cholate (0.25 g/L), sodium chenodeoxycholate (0.25 g/L), and Tween 80 (2 mL/L) in ddH_2_O. The base medium was sterilized by autoclaving at 121 °C for 15 minutes. Hemin (0.01 g/L) and vitamin K1 (10 µg/L) were sterilized by passing through a 0.22 µm filter (Cytiva, Cat: 9913-2502), and mucin (4.0 g/L) was weighed under sterile conditions; these components were then added to the base medium on the day of culturing.

PP was prepared according to Geerlings et al.[26] Briefly, PP base medium was prepared by combining pea peptone (32 g/L; OrganoTechnie), L-threonine (4 g/L), sodium bicarbonate (4 g/L), potassium dihydrogen phosphate (0.4 g/L), disodium hydrogen phosphate (0.669 g/L), ammonium chloride (0.3 g/L), sodium chloride (0.3 g/L), and magnesium chloride hexahydrate (0.1 g/L) in ddH_2_O, followed by sterilization at 121 °C for 15 minutes. The vitamin solution contained biotin (0.02 g/L), niacin (0.2 g/L;), pyridoxine (0.5 g/L), riboflavin (0.1 g/L), thiamine (0.2 g/L), cyanocobalamin (0.1 g/L), p-aminobenzoic acid (0.1 g/L), and pantothenic acid (0.1 g/L), and was sterile filtered prior to use. The glucose/N-acetylglucosamine supplement was prepared as a sterile filtered stock solution containing glucose and N-acetylglucosamine at a 3:1 molar ratio. Both the vitamin solution (final concentration of 1% v/v) and the glucose/N-acetylglucosamine supplement (final total concentration of 75 mM) were added to the autoclaved PP medium on the day of inoculation.

For all media, L-cysteine (0.5 g/L) was used as a reducing agent and added under anaerobic conditions prior to use. All reagents were purchased from Sigma-Aldrich, unless otherwise indicated.

### In vitro microbiome culturing and sample collection

Frozen microbiome biobank slurry was thawed in a 37°C water bath quickly with shaking, and immediately transferred into the anaerobic chamber. Fecal slurry was initially inoculated at an inoculation rate of 0.25% into the respective culture media in a 96 deep-well plate and incubated at 37°C with shaking at 500 rpm for 48 hours. Each donor-media condition was performed in four replicate wells. Following 48-hour incubation, an aliquot of each culture was inoculated into a new 96 deep-well plate containing the corresponding culture medium at a passaging inoculation rate of 0.5% and cultured for 48 hours. This passaging process was repeated sequentially for a total of 25 passages. At each passage, the remaining microbiome cultures were collected, centrifuged at 4000 rpm for 30 minutes at 4 °C to pellet microbial cells. The resulting microbial cell pellets were re-suspended with cold phosphate-buffered saline (PBS) and centrifuged again to wash the pellets. The microbial pellets were then stored in -80 °C until further metaproteomic processing.

A baseline uncultured sample, namely P0, was collected as well for each condition and microbiome by mixing 20% fecal slurry with respective culture medium to reach a final 2% fecal slurry to mimic the medium impacts on metaproteomics measurement. These samples were processed the same way as described above and the resulting microbial pellets were stored at -80 °C until further analysis.

### Optical density and pH measurement

Microbial growth was monitored by measuring optical density at 600 nm (OD600) with BioTek Epoch 2 microplate spectrophotometer (Agilent). Culture samples were diluted 1:1 with PBS prior to measurement. Corresponding media blanks were included for background correction.

The supernatant of the cultures was collected for pH measurement using an Accumet AB315 pH meter (Fisher Scientific). Where necessary, aliquots from replicate cultures were pooled to obtain sufficient volume for accurate and consistent measurements.

### Protein extraction and quantification

Microbial pellets were lysed with a lysis buffer composed of 4 M urea, 4% sodium dodecyl sulfate (SDS), and Halt™ protease inhibitor (10µL/1mL; ThermoFisher) in 100 mM Tris-HCl with ultrasonication in a cup-horn ultrasonicator Q700 maintained at 8 °C [50]. Sonication was performed using a 20-minute cycle (10 seconds on, 10 seconds off) at 50% amplitude. Following lysis, samples were centrifuged at 4000 rpm for 20 minutes at 8 °C, and the supernatant was transferred to a new plate for protein quantification using the Pierce™ BCA Protein Assay Kit (ThermoScientific) following the manufacturer’s instructions.

### Trypsin digestion and peptide clean-up

Trypsin digestion was performed using the single-pot, solid-phase-enhanced sample preparation (SP3) method as described previously with minor modifications [51]. Briefly, 10 µg of protein per sample aliquoted in a PCR plate was first reduced with dithiothreitol (DTT) and alkylated with iodoacetamide (IAA). Magnetic beads were prepared by combining equal amount of Sera-Mag carboxylate-modified speed beads A and B (Thermo Scientific), and added to the protein lysate at a 1:10 protein-to-bead mass ratio. Protein binding was induced by adding acetonitrile to a final concentration of around 70% (v/v), followed by incubation on a thermomixer at 1000 rpm for 20 minutes at room temperature to facilitate protein-bead binding. After binding, magnetic beads with proteins were immobilized using a magnetic rack, and the supernatant was removed. The beads were then washed 3-4 times with 80% (v/v) acetonitrile and one final wash with 100% acetonitrile. The proteins were resuspended in 20 µL of 50 mM ammonium bicarbonate (ABC) and briefly sonicated for homogenization and digested with 0.4 µg trypsin at 37 °C overnight with shaking at 1000 rpm.

Following digestion, samples were briefly sonicated and placed on a magnetic rack to settle down the beads. The supernatant was then collected and transferred to a new PCR plate. Samples were acidified with 10% (v/v) formic acid (FA) and desalted with Sep-Pak tC18 µElution Plate (Waters) according to manufacturer’s instructions. Briefly, the C18 beads were activated with 100% acetonitrile and equilibrated with 0.1% FA prior to sample loading; the beads with samples were washed with 0.1% FA and eluted with 50% acetonitrile / 0.1% FA. The eluents were then dried with a centrivap, and resuspended in 0.1% FA for LC-MS analysis.

### Liquid chromatography-mass spectrometry analysis

LC-MS analysis was performed using an Orbitrap Astral MS (ThermoFisher Scientific, San Jose, CA) in a data-independent acquisition (DIA) mode. Peptides were separated using the Vanquish Neo UHPLC with 75 μm I.D. × 150 mm fused silica analytical column packed in-house with 3μm ReproSil-Pur C18 beads (100 Å; Dr. Maisch GmbH, Ammerbuch, Germany). The solvent system consists of buffer A of 0.1% FA in water, and buffer B of 0.1% FA in 80% acetonitrile. LC separation was at a flow rate of 500nL/minute, and the gradient of 5-45% buffer B in 13.5 minutes.

The UHPLC system was coupled with the Orbitrap Astral mass spectrometer using a nano-electrospray interface operated in positive ion mode. The spray voltage used was 2.6 kV and the temperature of the heated capillary was 300 °C. DIA mode was used and the scan ranged from 380 to 980 m/z for both MS and DIA. The DIA window is set to 2 Dalton, with HCD energy of 25%. All data was recorded with Xcalibur software and exported as .RAW format for further bioinformatics processing.

### Metaproteomics data processing

Microbial peptide and protein identification, quantification and downstream taxonomic and functional annotation were performed using MetaPilot version 1.0 [24]. The Unified Human Gastrointestinal Genome v2.0 (UHGG v2.0) database which contained 4744 species representative reference genomes was downloaded from MGnify and used for database search [52, 53]. To build a spectral-library enabling data processing for large scale DIA-MS metaproteomics dataset in this study, 110 representative raw files samples were selected for a first MetaPilot database search to generate a reduced identified peptide sequence database (.fasta). The representative files were selected from six representative passages (P0, P1, P3, P7, P15, and P20), with one raw file selected for each donor within each culture medium, resulting in 108 raw files. Two QC raw files were also included, yielding a total of 110 raw files. Briefly, the DDA-independent Marker4 DIA workflow of MetaPilot was used to search the 110 representative raw files against entire UHGG genome database. In total 1,646,472 peptide sequences from 844 genomes were identified and subsequently used for database search of the entire dataset (1008 microbiome and 9 QC). For this full search, the seeded DIA search workflow (direct search mode) in MetaPilot was used. In both searches, DIA-NN 2.3 was used for identification and quantification with the following parameters [54]: match-between-run (MBR) was disabled; peptide length 7-30; charge 2-3; MS1 range 350-1200; MS2 range 200-1800, and MS1 and MS2 accuracy of 4ppm and 10ppm, respectively; false discovery rate (FDR) was set as default (1% at both precursor and protein levels).

Identified peptides were traced back to UHGG database genome information, which was then used to identify and quantify genomes. The genome annotation information and the quantification table were used for taxonomic analysis at different taxonomic levels. Similarly, functional annotation of identified proteins was obtained directly from the UHGG database and used for functional analysis.

For virulence factor (VF) analysis, VFDB was downloaded from https://www.mgc.ac.cn/VFs/ [45]. The full VFDB database was concatenated with all identified microbial peptides in MetaPilot database search. DIA-NN 2.3 was used for a library-free search against this database for the entire dataset. VF category information was obtained directly from the fasta header. For antimicrobial resistant protein analysis, the Comprehensive Antibiotic Resistance Database (CARD; https://card.mcmaster.ca/) [55] was concatenated with all identified microbial peptides, and DIA-NN 2.3 was used for a library-free search for all 1017 samples.

For phage proteome analysis, the UHGG v2.0 database was combined directly with the Gut Phage Database (GPD; https://www.sanger.ac.uk/data/gut-phage-database/) [36], which contains 142,809 phage genomes. The same set of 110 representative samples was used in the first search to generate a phage-inclusive spectral library using DIA-NN 2.3 with an InfinDIA mode. This library was subsequently applied to library search across all 1017 samples using DIA-NN as well. Taxonomic assignment of phage proteins was obtained directly from the GPD database.

### Taxonomic and diversity analysis

Taxonomic analysis was performed with genome abundances. Taxonomic annotation of genomes was extracted directly from the UHGG database and used for calculation of relative abundances at different taxonomic levels. The database used in this metaproteomic analysis includes 4744 species-representative, and thereby the genomes identified or quantified here are equivalent to species. Taxonomic composition was visualized using stacked bar plots, and alpha diversity was assessed using Shannon diversity using the species abundance table. Bray-Curtis and UniFrac dissimilarities in species composition were calculated using R package XX, and the phylogenetic tree data was downloaded from the MGnify database. Phylogenetic tree display of identified species was performed using MetaTree (https://byemaxx.github.io/MetaTree/) [56].

Species retention analysis was performed using a three-level filtering strategy. Briefly, a species needs to have at least three unique peptides identified across the entire dataset; at each sample level, a species was considered present only if it reached a minimum relative abundance of 0.1%, and at the passage level, a species was considered present in the passage only if it was detected in at least two replicates.

### Operational taxon-function (OTF) analysis

MetaX version 1.0 was used to link functions to taxa to generate OTF [31]. MetaX uses identified peptide identification results (final_peptides.tsv or pr_matrix.tsv) to directly generate OTFs based on pre-built taxa and function database in MetaX. The lowest common ancestor (LCA) threshold was set as 1 and genomes were filtered with a minimum of 3 distinct peptides to build OTF from peptide tables. In this study, OTF analysis was performed at the genus level to balance the taxonomic resolution and information retention (proportion of peptides used). Genus-KO and genus-CAZy OTF tables were exported from MetaX for further statistical analysis and visualization.

Statistical analysis and visualization

PCA was used to visualize overall variation in the dataset. Intensity data of both proteins, functions and taxa were log-transformed and filtered to retain features present in at least 80% of samples. Missing values were imputed using a k-nearest neighbors (kNN) approach, unless otherwise indicated. Distance analysis was performed based on Euclidean distances in the principal component (PC) space to quantify compositional deviation from uncultured baseline (P0) microbiomes. For each donor, a baseline centroid was defined as the mean PC coordinates of P0 samples, and the Euclidean distance between each sample and its corresponding donor-specific P0 centroid was computed, providing a measure of compositional divergence across passages and media conditions. Permutational Multivariate Analysis of Variance (PERMANOVA) was used to evaluate the contribution and significance of different factors, including medium, donor and their interactions, to the observed variations in PCA analysis.

Supervised PLS-DA was used to identify differentially abundant functions with a threshold of variable importance in the projection (VIP) ≥1. Two group comparisons were performed using Wilcoxon rank-sum tests and fold changes (FC). A feature with a BH-adjusted p <0.05 and FC ≥ 2 was considered significant between groups. The comparison results were visualized using a Volcano plot or heatmap. All data analysis and visualization were performed using R (version 4.5.1) unless otherwise indicated. AI tools (ChatGPT, Gemini and Cursor) were used to assist with R code debugging and refinement. All AI-assisted code was reviewed, modified as needed, validated by the authors and executed locally using RStudio (version 2024.09.1 Build 394). The authors take full responsibility for the analysis and interpretation of the results.

## Data and code availability

The mass spectrometry data have been deposited to the ProteomeXchange Consortium (http://www.proteomexchange.org). Any additional information required to reanalyze the data reported in this paper is available from the lead contact.

## Acknowledgments

This study was supported by the Health Canada and Phase VIII Genomics R&D Initiative (GRDI) grant of the Government of Canada. We gratefully acknowledge Drs Karima Benkhedda, Eric Deneault and Jessie Lavoie from Health Canada for their critical comments and edits to this manuscript.

## Author contributions

Study design and supervision: XZ, DF, XL, VD; live microbiome biobank sample collection: JM, DF; experiment, sample collection, processing and data acquisition: AW, XZ, ZN, EF; data analysis and visualization: AW, XZ, KC, QW; manuscript drafting: XZ, AW; administrative, technical, or material support: MC, EF, JM, DF. All authors revised and edited the manuscript.

## References

1. O’Hara, A.M. and F. Shanahan, The gut flora as a forgotten organ. EMBO Rep, 2006. 7(7): p. 688–93.

2. Bi, Y., et al., The gut microbiome organ. Imeta, 2026: p. e70144.

3. Carding, S., et al., Dysbiosis of the gut microbiota in disease. Microb Ecol Health Dis, 2015. 26: p. 26191.

4. Lin, X., et al., Gut-X axis. Imeta, 2025. 4(1): p. e270.

5. Feuerstadt, P., et al., SER-109, an Oral Microbiome Therapy for Recurrent Clostridioides difficile Infection. N Engl J Med, 2022. 386(3): p. 220–229.

6. McGovern, B.H., et al., SER-109, an Investigational Microbiome Drug to Reduce Recurrence After Clostridioides difficile Infection: Lessons Learned From a Phase 2 Trial. Clin Infect Dis, 2021. 72(12): p. 2132–2140.

7. Sims, M.D., et al., Safety and Tolerability of SER-109 as an Investigational Microbiome Therapeutic in Adults With Recurrent Clostridioides difficile Infection: A Phase 3, Open-Label, Single-Arm Trial. JAMA Netw Open, 2023. 6(2): p. e2255758.

8. Khanna, S., et al., Efficacy and Safety of RBX2660 in PUNCH CD3, a Phase III, Randomized, Double-Blind, Placebo-Controlled Trial with a Bayesian Primary Analysis for the Prevention of Recurrent Clostridioides difficile Infection. Drugs, 2022. 82(15): p. 1527–1538.

9. Fekete, E.E., D. Figeys, and X. Zhang, Microbiota-directed biotherapeutics: considerations for quality and functional assessment. Gut Microbes, 2023. 15(1): p. 2186671.

10. Li, L., et al., RapidAIM 2.0: a high-throughput assay to study functional response of human gut microbiome to xenobiotics. Microbiome Res Rep, 2024. 3(2): p. 26.

11. Li, L., et al., RapidAIM: a culture- and metaproteomics-based Rapid Assay of Individual Microbiome responses to drugs. Microbiome, 2020. 8(1): p. 33.

12. Li, L., et al., Systematic metaproteomics mapping reveals functional and ecological landscapes of Ex vivo human gut microbiota responses to therapeutic drugs. Nat Commun, 2025. 16(1): p. 9383.

13. Li, L., et al., A functional ecological network based on metaproteomics responses of individual gut microbiomes to resistant starches. Comput Struct Biotechnol J, 2020. 18: p. 3833–3842.

14. Dobranowski, P., et al., Individual variability shapes ex vivo responses to resistant starch in inflammatory bowel disease derived microbiomes. NPJ Biofilms Microbiomes, 2026. 12(1).

15. Li, L., et al., Berberine and its structural analogs have differing effects on functional profiles of individual gut microbiomes. Gut Microbes, 2020. 11(5): p. 1348–1361.

16. Van de Wiele, T., et al., The simulator of the human intestinal microbial ecosystem (SHIME®), in The Impact of Food Bioactives on Health: in vitro and ex vivo models, K. Verhoeckx, et al., Editors. 2015, Springer. p. 305-317.

17. Zhu, W., et al., Simulator of the Human Intestinal Microbial Ecosystem (SHIME((R))): Current Developments, Applications, and Future Prospects. Pharmaceuticals (Basel), 2024. 17(12).

18. Cheng, A.G., et al., Design, construction, and in vivo augmentation of a complex gut microbiome. Cell, 2022. 185(19): p. 3617–3636 e19.

19. Aranda-Diaz, A., et al., Establishment and characterization of stable, diverse, fecal-derived in vitro microbial communities that model the intestinal microbiota. Cell Host Microbe, 2022. 30(2): p. 260–272 e5.

20. Aranda-Diaz, A., et al., Assembly of stool-derived bacterial communities follows "early-bird" resource utilization dynamics. Cell Syst, 2025. 16(4): p. 101240.

21. Zhang, X. and D. Figeys, Perspective and Guidelines for Metaproteomics in Microbiome Studies. J Proteome Res, 2019. 18(6): p. 2370–2380.

22. Kleiner, M., Metaproteomics: Much More than Measuring Gene Expression in Microbial Communities. mSystems, 2019. 4(3).

23. Zhang, X., et al., Clinical Microbiome Analysis by Mass Spectrometry-Based Metaproteomics. Annu Rev Anal Chem (Palo Alto Calif), 2025. 18(1): p. 149–172.

24. Cheng, K. and D. Figeys, MetaPilot enables adaptive genome-aware DDA and DIA metaproteomics using microbial genome catalogues. bioRxiv, 2026: p. 2026.06.12.728088.

25. Dumas, T., et al., The astounding exhaustiveness and speed of the Astral mass analyzer for highly complex samples is a quantum leap in the functional analysis of microbiomes. Microbiome, 2024. 12(1): p. 46.

26. Geerlings, S.Y., et al., Omics-based analysis of Akkermansia muciniphila cultivation in food-grade media. Microbiome Res Rep, 2024. 3(3): p. 36.

27. Guan, N. and L. Liu, Microbial response to acid stress: mechanisms and applications. Appl Microbiol Biotechnol, 2020. 104(1): p. 51–65.

28. Imlay, J.A., The molecular mechanisms and physiological consequences of oxidative stress: lessons from a model bacterium. Nat Rev Microbiol, 2013. 11(7): p. 443–54.

29. Tripathi, P., et al., ClpB is an essential stress regulator of Mycobacterium tuberculosis and endows survival advantage to dormant bacilli. Int J Med Microbiol, 2020. 310(3): p. 151402.

30. Song, C., et al., Intestinal mucus components and secretion mechanisms: what we do and do not know. Exp Mol Med, 2023. 55(4): p. 681–691.

31. Wu, Q., et al., Operational Taxon-Function Framework in MetaX: Unveiling Taxonomic and Functional Associations in Metaproteomics. Anal Chem, 2025. 97(18): p. 9739–9747.

32. Ducarmon, Q.R., et al., Cayman enables large-scale analysis of gut microbiome carbohydrate-active enzyme repertoires. Nat Microbiol, 2026. 11(6): p. 1739–1753.

33. Blakeley-Ruiz, J.A. and M. Kleiner, Considerations for constructing a protein sequence database for metaproteomics. Comput Struct Biotechnol J, 2022. 20: p. 937–952.

34. Van Den Bossche, T., et al., The microbiologist’s guide to metaproteomics. Imeta, 2025. 4(3): p. e70031.

35. Van Den Bossche, T., et al., The need for standardization and improved open (meta)data practices in metaproteomics. Microbiome, 2026. 14(1).

36. Camarillo-Guerrero, L.F., et al., Massive expansion of human gut bacteriophage diversity. Cell, 2021. 184(4): p. 1098–1109 e9.

37. Shkoporov, A.N., et al., The Human Gut Virome Is Highly Diverse, Stable, and Individual Specific. Cell Host Microbe, 2019. 26(4): p. 527–541 e5.

38. Li, L., et al., Evaluating in Vitro Culture Medium of Gut Microbiome with Orthogonal Experimental Design and a Metaproteomics Approach. J Proteome Res, 2018. 17(1): p. 154–163.

39. Li, L., et al., An in vitro model maintaining taxon-specific functional activities of the gut microbiome. Nat Commun, 2019. 10(1): p. 4146.

40. Tailford, L.E., et al., Mucin glycan foraging in the human gut microbiome. Front Genet, 2015. 6: p. 81.

41. Martens, E.C., H.C. Chiang, and J.I. Gordon, Mucosal glycan foraging enhances fitness and transmission of a saccharolytic human gut bacterial symbiont. Cell Host Microbe, 2008. 4(5): p. 447–57.

42. Koropatkin, N.M., E.A. Cameron, and E.C. Martens, How glycan metabolism shapes the human gut microbiota. Nat Rev Microbiol, 2012. 10(5): p. 323–35.

43. Wu, E., et al., Data-independent acquisition in metaproteomics. Expert Rev Proteomics, 2024. 21(7-8): p. 271–280.

44. Wang, A., et al., Assessing fecal metaproteomics workflow and small protein recovery using DDA and DIA PASEF mass spectrometry. Microbiome Res Rep, 2024. 3(3): p. 39.

45. Zhou, S., et al., VFDB 2025: an integrated resource for exploring anti-virulence compounds. Nucleic Acids Res, 2025. 53(D1): p. D871–D877.

46. Diebold, P.J., et al., Clinically relevant antibiotic resistance genes are linked to a limited set of taxa within gut microbiome worldwide. Nat Commun, 2023. 14(1): p. 7366.

47. Dong, W., et al., An expanded database and analytical toolkit for identifying bacterial virulence factors and their associations with chronic diseases. Nat Commun, 2024. 15(1): p. 8084.

48. Van Rossum, T., et al., Diversity within species: interpreting strains in microbiomes. Nat Rev Microbiol, 2020. 18(9): p. 491–506.

49. Zhang, X., et al., Evaluating live microbiota biobanking using an ex vivo microbiome assay and metaproteomics. Gut Microbes, 2022. 14(1): p. 2035658.

50. Zhang, X., et al., Assessing the impact of protein extraction methods for human gut metaproteomics. J Proteomics, 2018. 180: p. 120–127.

51. Hughes, C.S., et al., Single-pot, solid-phase-enhanced sample preparation for proteomics experiments. Nat Protoc, 2019. 14(1): p. 68–85.

52. Almeida, A., et al., A unified catalog of 204,938 reference genomes from the human gut microbiome. Nat Biotechnol, 2021. 39(1): p. 105–114.

53. Richardson, L., et al., MGnify: the microbiome sequence data analysis resource in 2023. Nucleic Acids Res, 2023. 51(D1): p. D753–D759.

54. Demichev, V., et al., DIA-NN: neural networks and interference correction enable deep proteome coverage in high throughput. Nat Methods, 2020. 17(1): p. 41–44.

55. Alcock, B.P., et al., CARD 2023: expanded curation, support for machine learning, and resistome prediction at the Comprehensive Antibiotic Resistance Database. Nucleic Acids Res, 2023. 51(D1): p. D690–D699.

56. Wu, Q., et al., MetaTree: an interactive web platform for aligned hierarchical data visualization and multi-group comparison. BMC Bioinformatics, 2026. 27(1).

