## Supplementary Figures for "Dissecting functional dynamics of fecal-derived *in vitro* stable microbial communities with metaproteomics"

^1^ Regulatory Research Division, Centre for Oncology, Biostatistics, Research and Radiopharmaceuticals, Biologic and Radiopharmaceutical Drugs Directorate, Health Products and Food Branch, Health Canada, Ottawa, Canada; ^2^ Department of Biochemistry, Microbiology and Immunology, Faculty of Medicine, University of Ottawa, Ottawa, Canada; ^3^ Quadram Institute Bioscience, Norwich Research Park, University of East Anglia, Norwich, Norfolk, United Kingdom; ^4^ Centre for Infection, Immunity and Inflammation, University of Ottawa, Ottawa, Canada; ^5^ School of Pharmaceutical Sciences, Faculty of Medicine, University of Ottawa, Ottawa, Canada.

* Correspondence to: (XZ), (DF), and (SL)

Supplementary Information:

### Supplementary Tables

**Table S1** Differentially abundant KOs between BHI- and RA-SICs. (.xlsx)

### Supplementary Figures

**Figure S1** Evaluation of quality control samples in metaproteomics experiment

**Figure S2** KEGG KO-based distance analysis for each individual

**Figure S3** PCA score plots based on quantified protein abundance profiles.

**Figure S4** Distance analysis based on the top protein PCA components.

**Figure S5** PERMANOVA of protein abundances for each passage.

**Figure S6** Phylum level composition of *in vitro* cultured microbiomes

**Figure S7** Family level composition of *in vitro* cultured microbiomes

**Figure S8** Species retention of *in vitro* culture microbiomes

**Figure S9** Differentially abundant OTFs matched to the top differential abundant KOs between BHI- and RA-SICs.

**Figure S10** Composition and functional dynamics of *Escherichia* and *Enterococcus* in PP-cultured microbiomes

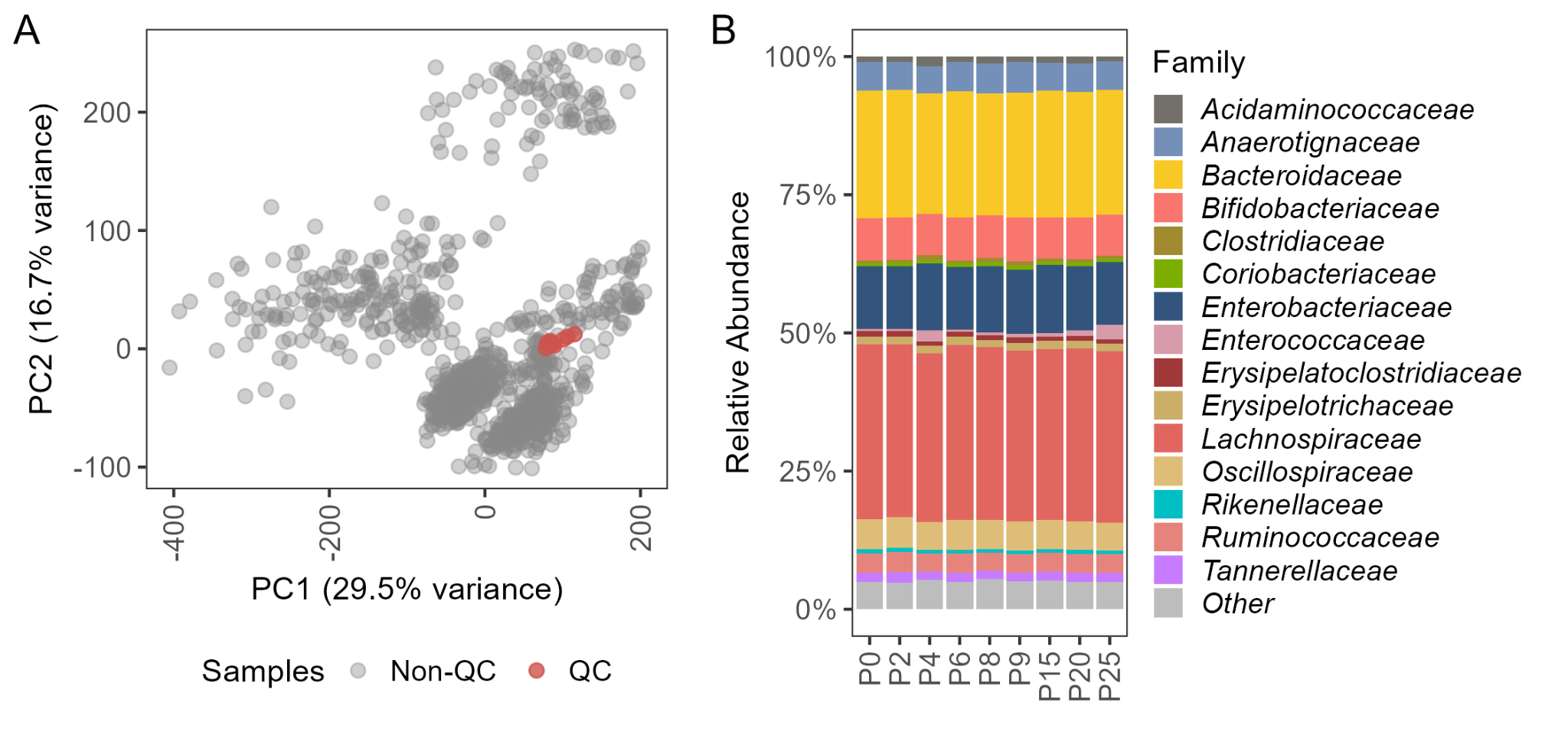

**Figure S1** Evaluation of quality control samples in metaproteomics experiment. *An identical quality control (QC) sample was run during different batches of LC-MS analysis together with the passage samples, and all resulting data were included for data analysis to evaluate the performance of LC-MS and metaproteomics workflow. (A) PCA score plot of quantified proteins showing QC samples are clustered closely together. (B) Family level composition of the QC samples showing consistent taxonomic composition throughout the experiment.*

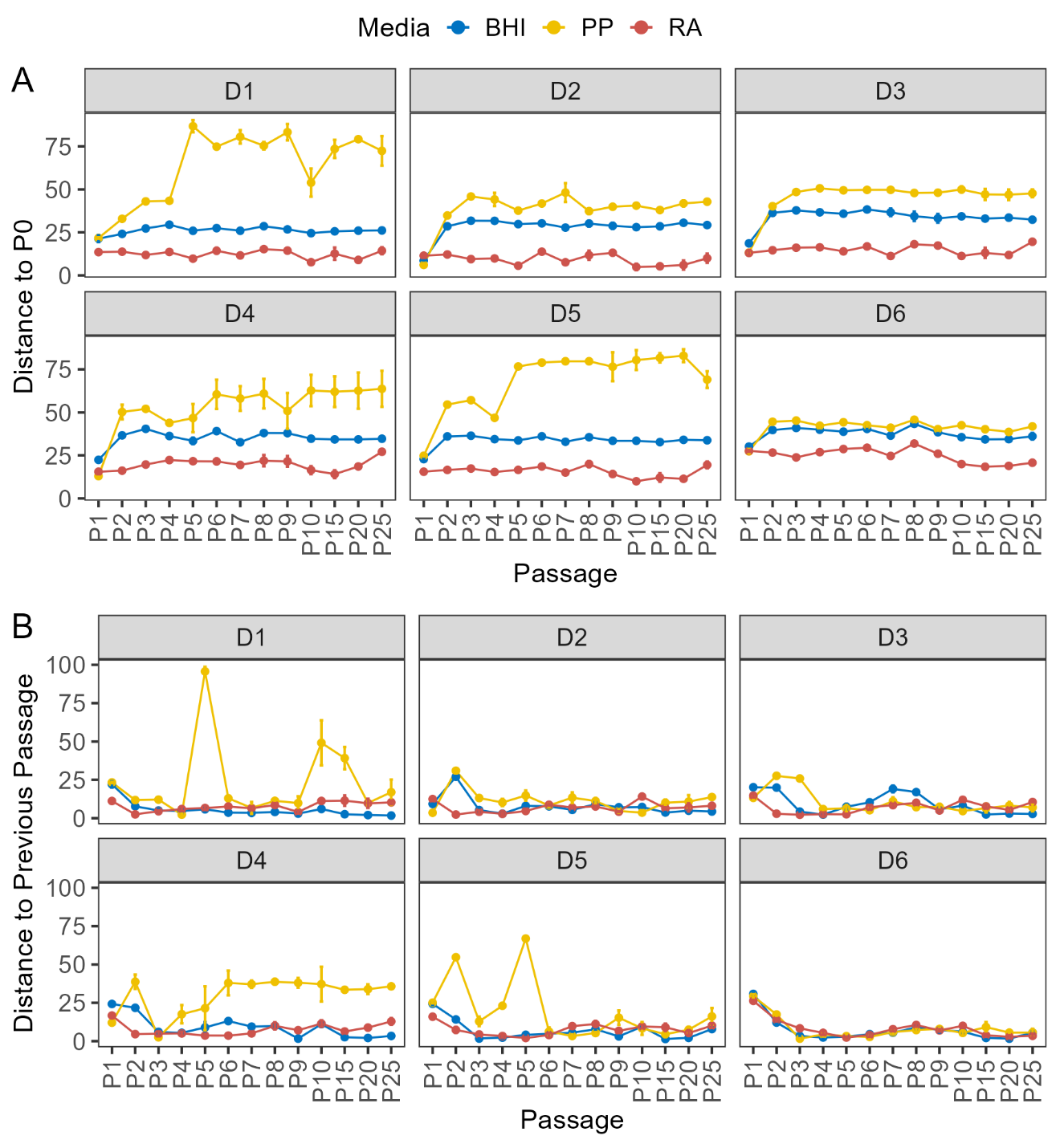

**Figure S2** KEGG KO-based distance analysis for each individual. *Distance analysis based on PCA score plots for each individual was performed with (A) Distance of each community to the corresponding P0 microbiome, and (B) distance to the previous passage.*

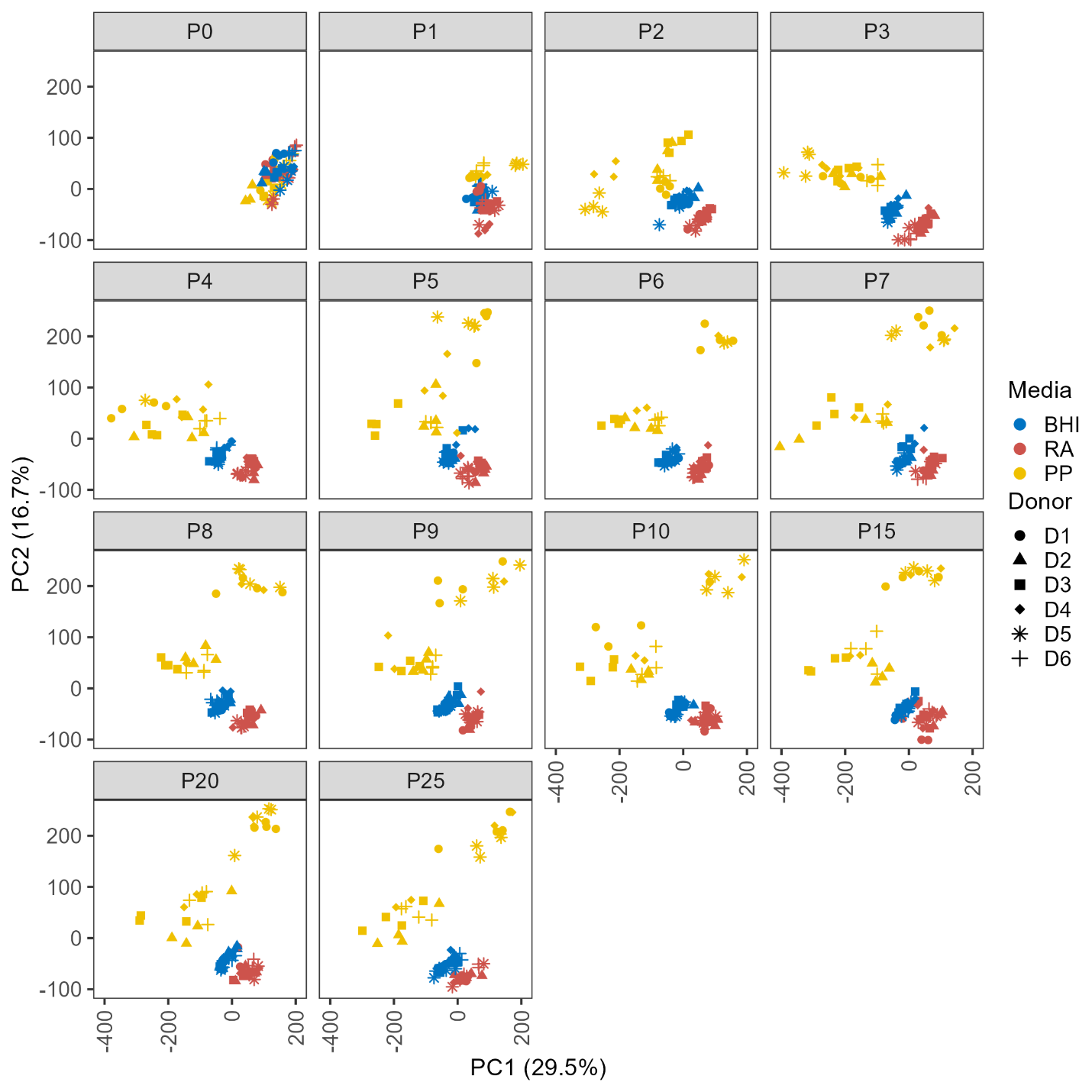

**Figure S3** PCA score plots based on quantified protein abundance profiles. Metaproteomic dynamics of *in vitro* cultured fecal microbiota. *PCA plot* was *faceted by passage, colors indicate culture media, and symbols indicate donor microbiomes.*

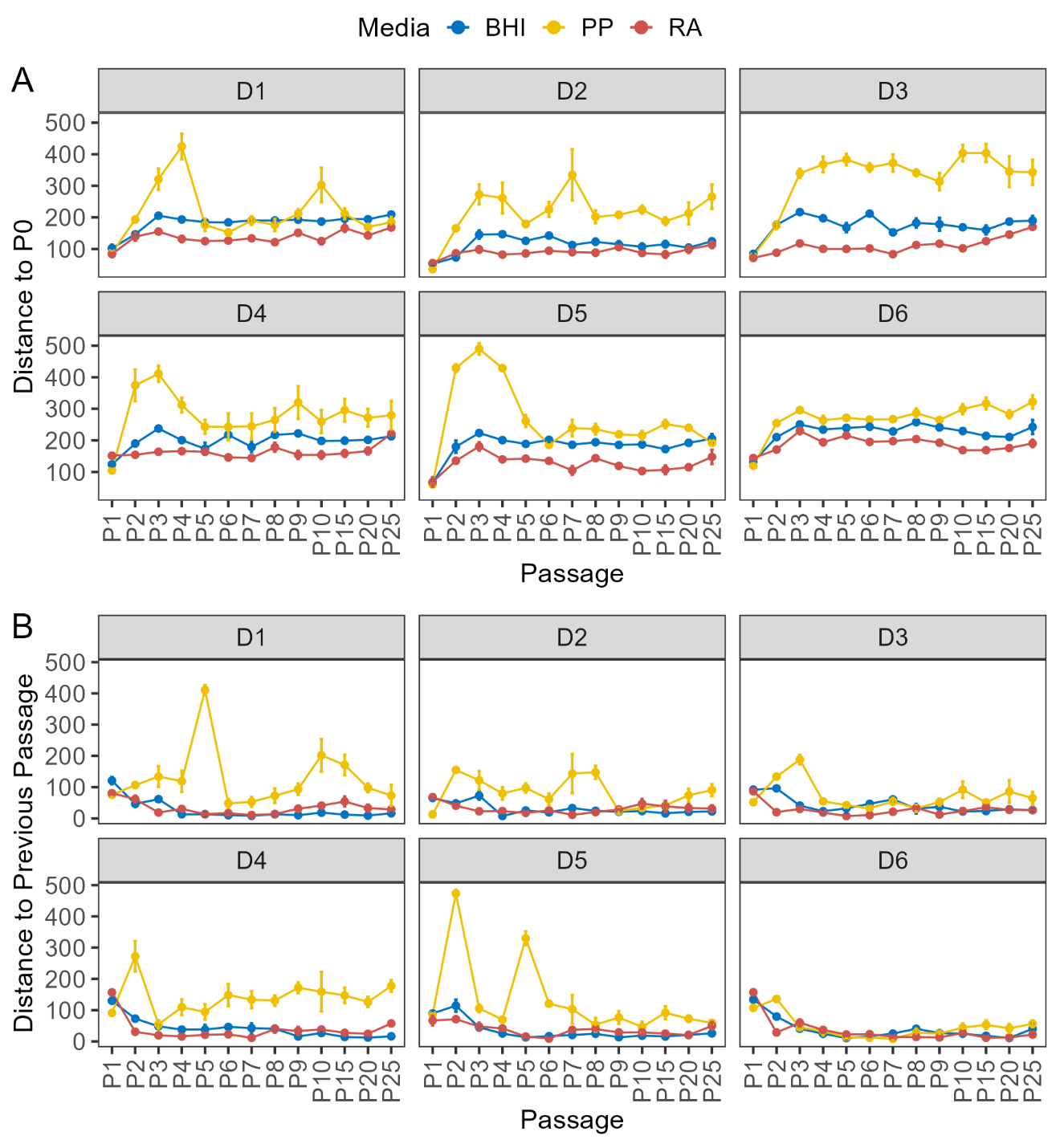

**Figure S4** Distance analysis based on protein PCA score plots. *Distance of each community to the corresponding P0 microbiome (A) and distance to the previous passage (B) were shown. Colors indicate different culture media.*

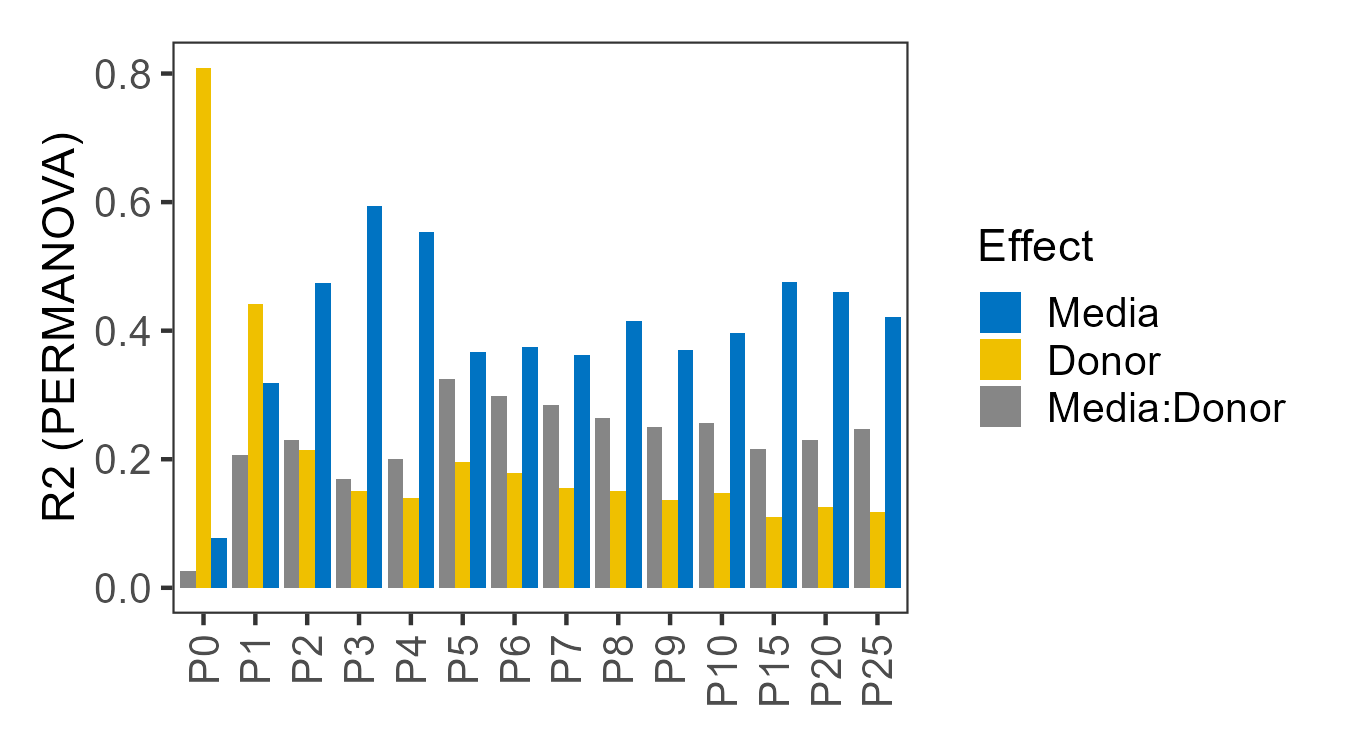

**Figure S5** PERMANOVA of protein abundances for each passage. *PCA components explaining 80% of the total variance were used for the analysis. Bars show the proportion of variance explained (R²) by culture medium, donor, and their interaction at each passage and colored according to the effect factor.*

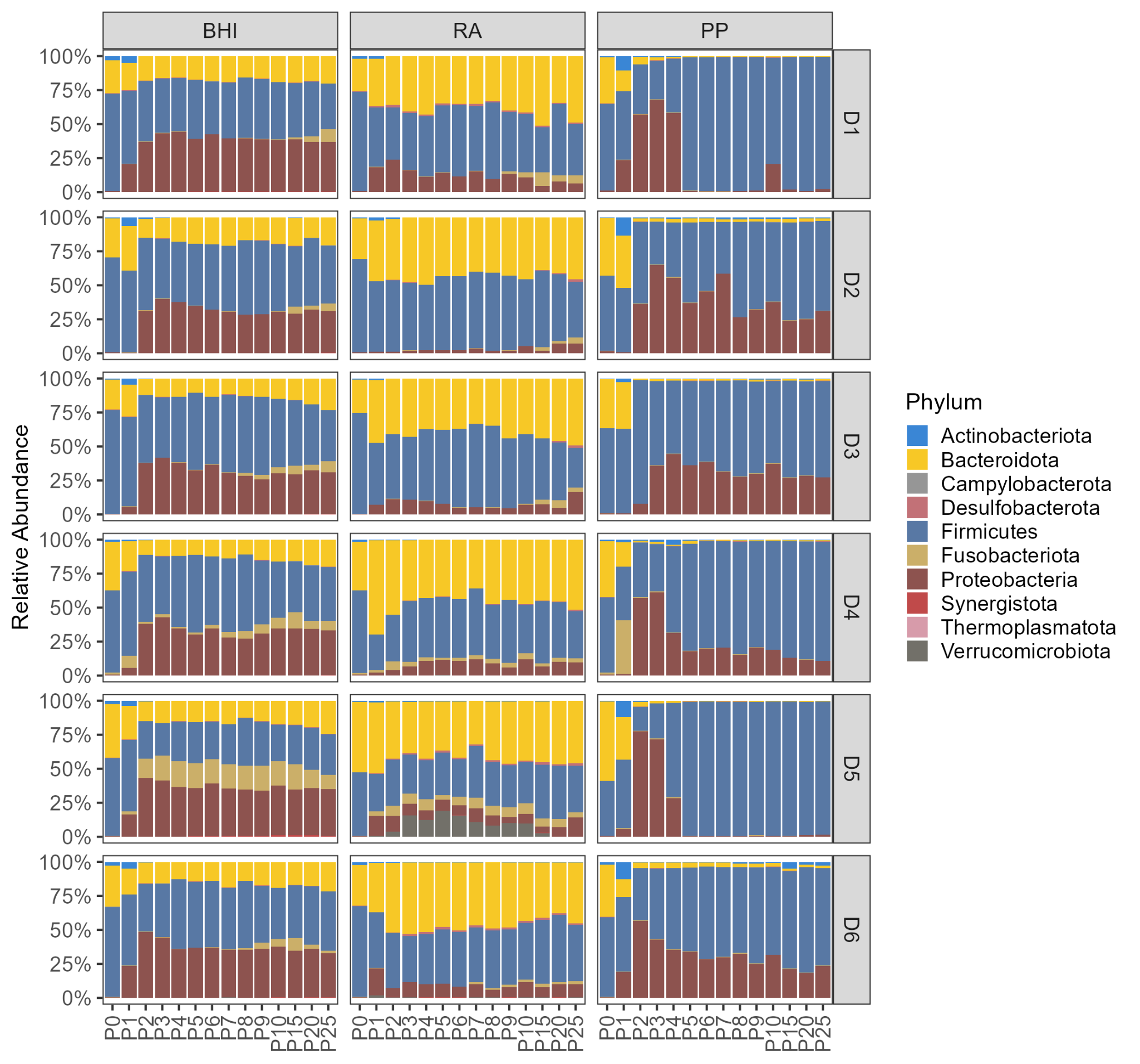

**Figure S6** Phylum level composition of *in vitro* cultured microbiomes. *Group means at each passage, faceted by culture media and donors were shown.*

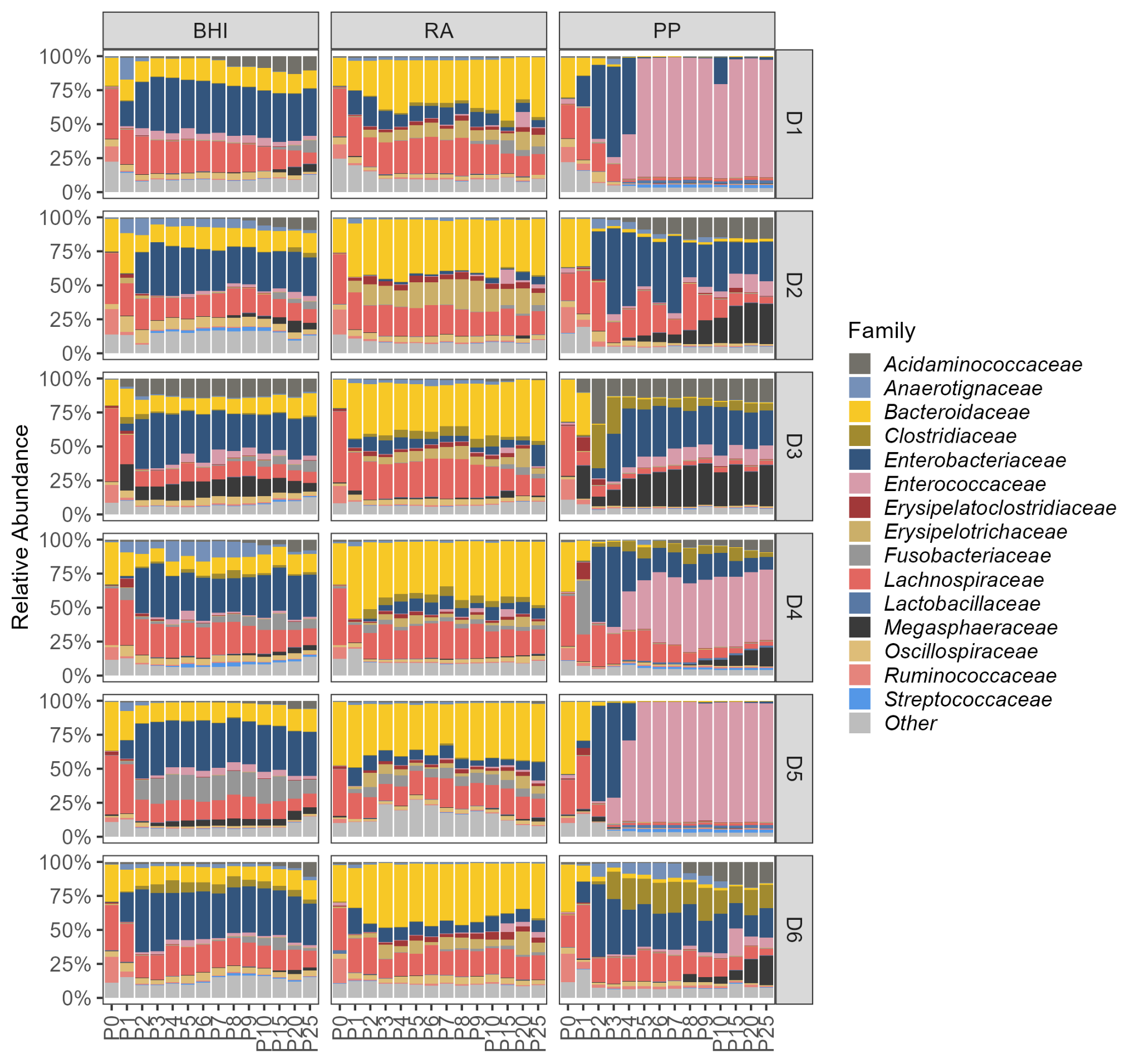

**Figure S7** Family level composition of *in vitro* cultured microbiomes. *Group means at each passage, faceted by culture media and donors were shown. Top 15 abundant families were shown with the rest families aggregated as Other.*

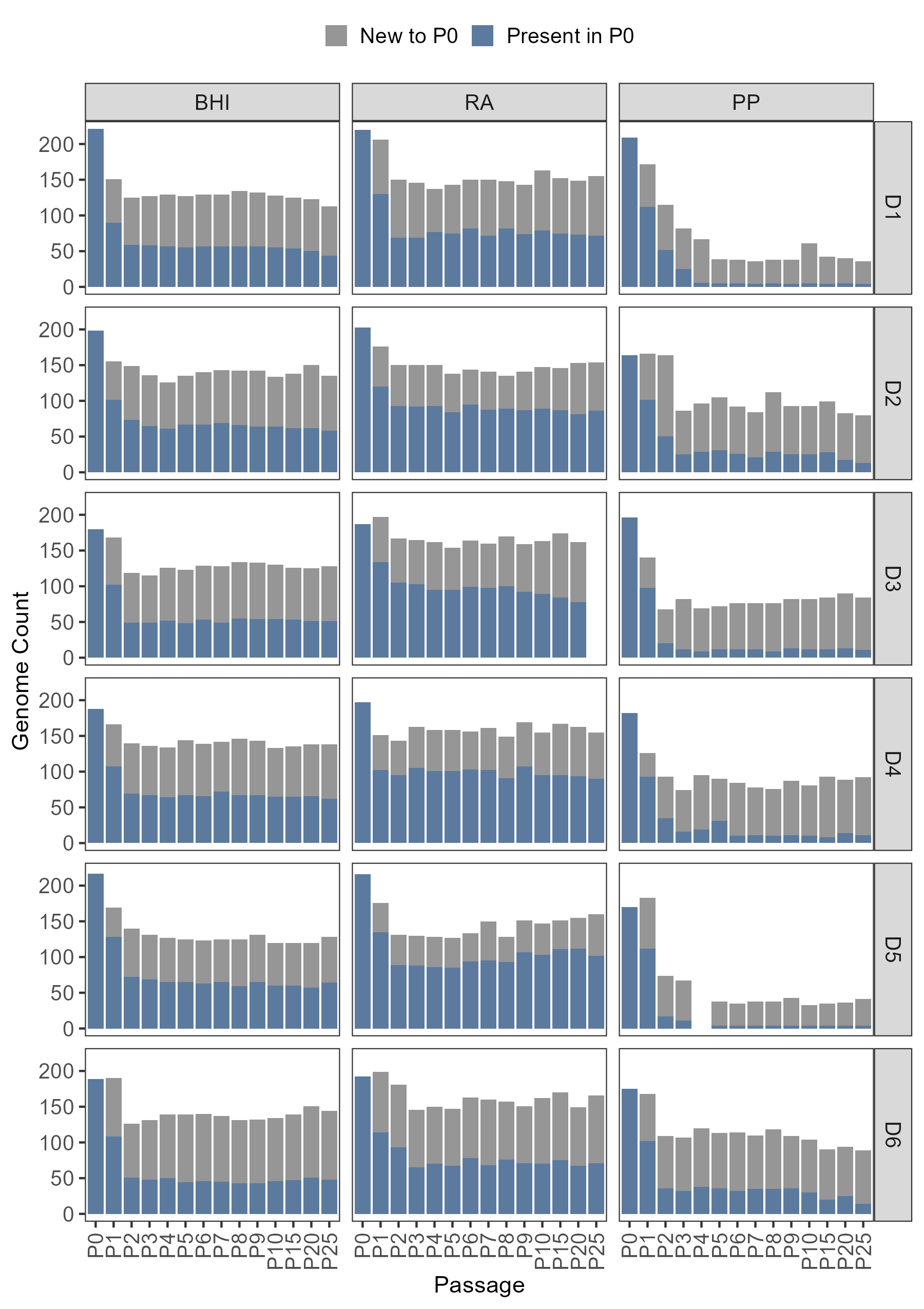

**Figure S8** Species retention of *in vitro* culture microbiomes. *A genome was considered present at a given passage, donor and medium pair if its abundance is >0.1% in at least two replicates from the same group. Stacked bars show genomes retained from P0 (dark blue) and genomes not detected at P0 (grey) for the following passages.*

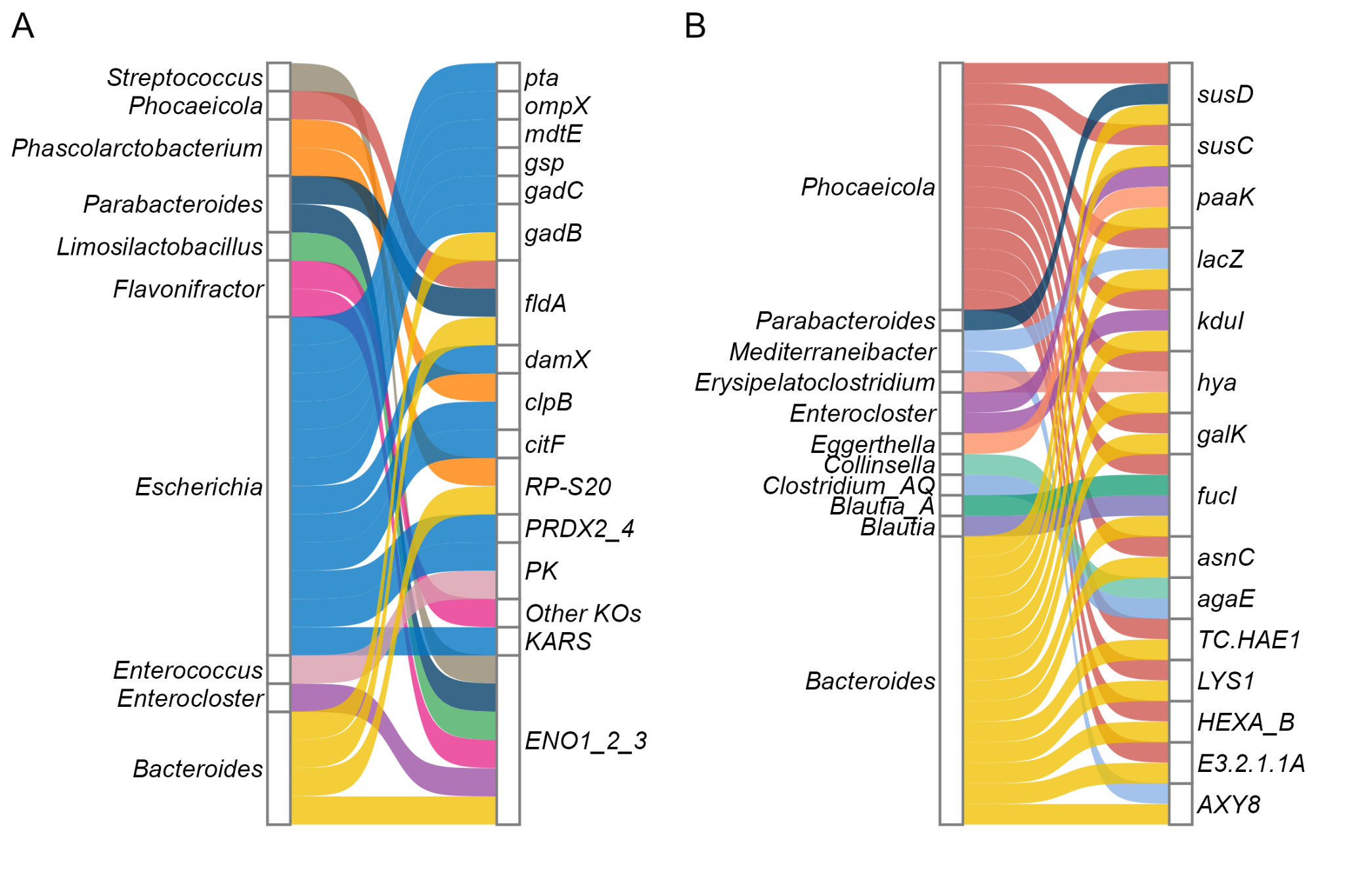

**Figure S9** Differentially abundant OTFs matched to the top differential abundant KOs between BHI- and RA-SICs. *Significantly different KO-genus OTFs that matched to the change direction of the identified top 50 differential abundant KOs upregulated in BHI-SIC (A) and RA-SIC / P0 microbiome (B). Ribbons are colored based on taxa and size corresponds to the number of significant OTFs.*

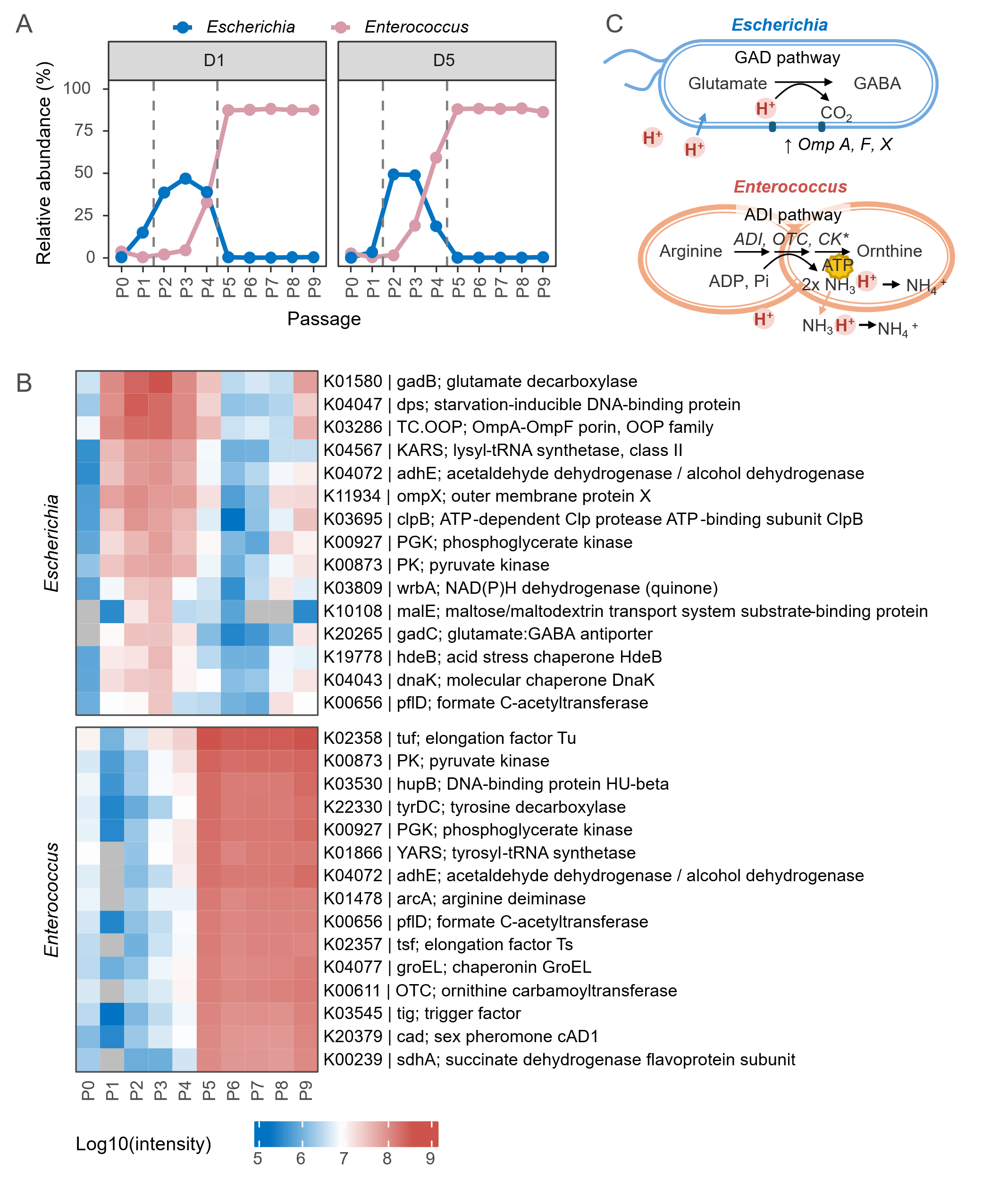

**Figure S10** Composition and functional dynamics of *Escherichia* and *Enterococcus* in PP-cultured microbiomes. *(A) Relative abundance of Escherichia and Enterococcus in early passages. (B) Heatmap of the top 15 abundant differentially OTFs linked to Escherichia (top) and Enterococcus (bottom), respectively. Genus-KO OTF data was used for comparisons of P2-P3 vs P0-P1 (top), and P4-P5 vs P0-P2, respectively, and top OTFs were selected based on the absolute intensity changes. (C) Diagram shows the acid resistance mechanisms utilized by Escherichia and Enterococcus supported by the metaproteomics data in this study.*
